# Distinct brain-wide presynaptic networks underlie the functional identity of individual cortical neurons

**DOI:** 10.1101/2023.05.25.542329

**Authors:** Ana R. Inácio, Ka Chun Lam, Yuan Zhao, Francisco Pereira, Charles R. Gerfen, Soohyun Lee

## Abstract

Neuronal connections provide the scaffolding for neuronal function. Revealing the connectivity of functionally identified individual neurons is necessary to understand how activity patterns emerge and support behavior. Yet, the brain-wide presynaptic wiring rules that lay the foundation for the functional selectivity of individual neurons remain largely unexplored. Cortical neurons, even in primary sensory cortex, are heterogeneous in their selectivity, not only to sensory stimuli but also to multiple aspects of behavior. Here, to investigate presynaptic connectivity rules underlying the selectivity of pyramidal neurons to behavioral state^1–12^ in primary somatosensory cortex (S1), we used two-photon calcium imaging, neuropharmacology, single-cell based monosynaptic input tracing, and optogenetics. We show that behavioral state-dependent neuronal activity patterns are stable over time. These are minimally affected by neuromodulatory inputs and are instead driven by glutamatergic inputs. Analysis of brain-wide presynaptic networks of individual neurons with distinct behavioral state-dependent activity profiles revealed characteristic patterns of anatomical input. While both behavioral state-related and unrelated neurons had a similar pattern of local inputs within S1, their long-range glutamatergic inputs differed. Individual cortical neurons, irrespective of their functional properties, received converging inputs from the main S1-projecting areas. Yet, neurons that tracked behavioral state received a smaller proportion of motor cortical inputs and a larger proportion of thalamic inputs. Optogenetic suppression of thalamic inputs reduced behavioral state-dependent activity in S1, but this activity was not externally driven. Our results revealed distinct long-range glutamatergic inputs as a substrate for preconfigured network dynamics associated with behavioral state.

## Introduction

Anatomical connectivity within and between brain areas governs the distinct activity patterns of individual neurons. In sensory cortical areas, the properties of local presynaptic connections, including their number, strength, and spatial arrangement, shape the selectivity of individual neurons to sensory stimuli^13–22^. Cortical neurons are, however, functionally highly heterogeneous in relation to various aspects of behavioral contexts and states, such as unexpected events, rewards, attentional demands, or spontaneous movements^1–12,23–27^. Even in sensory cortical areas, neurons show highly dynamic activity in the absence of external sensory stimuli^1,7,11^. These behavioral contextual and state signals are thought to be conveyed by long-range projections, including neuromodulatory and glutamatergic afferents from multiple brain areas^28–36^.

One way to establish functional heterogeneity is that a specific set of inputs projects to a subset of neurons, whether the inputs are neuromodulatory or long-range and local glutamatergic and GABAergic, or all the above (Fig. 1a). Alternatively, presynaptic inputs may be random and highly plastic (Fig. 1a). While the mesoscale level of connectivity within and between brain areas has been mapped^37^, linking connectivity rules with the functional identity of individual neurons remains challenging. To address this question, we investigated the architecture of brain-wide presynaptic connectivity in the context of spontaneous movement-sensitive neurons in the primary somatosensory cortex. Spontaneous movements, including whisking and locomotion, are a component of innate behavior. These spontaneous movements are robustly represented in a subset of neurons in a wide range of brain areas^9–11^. However, how the functional specificity of these neurons arises is not clear.

**Fig. 1:**
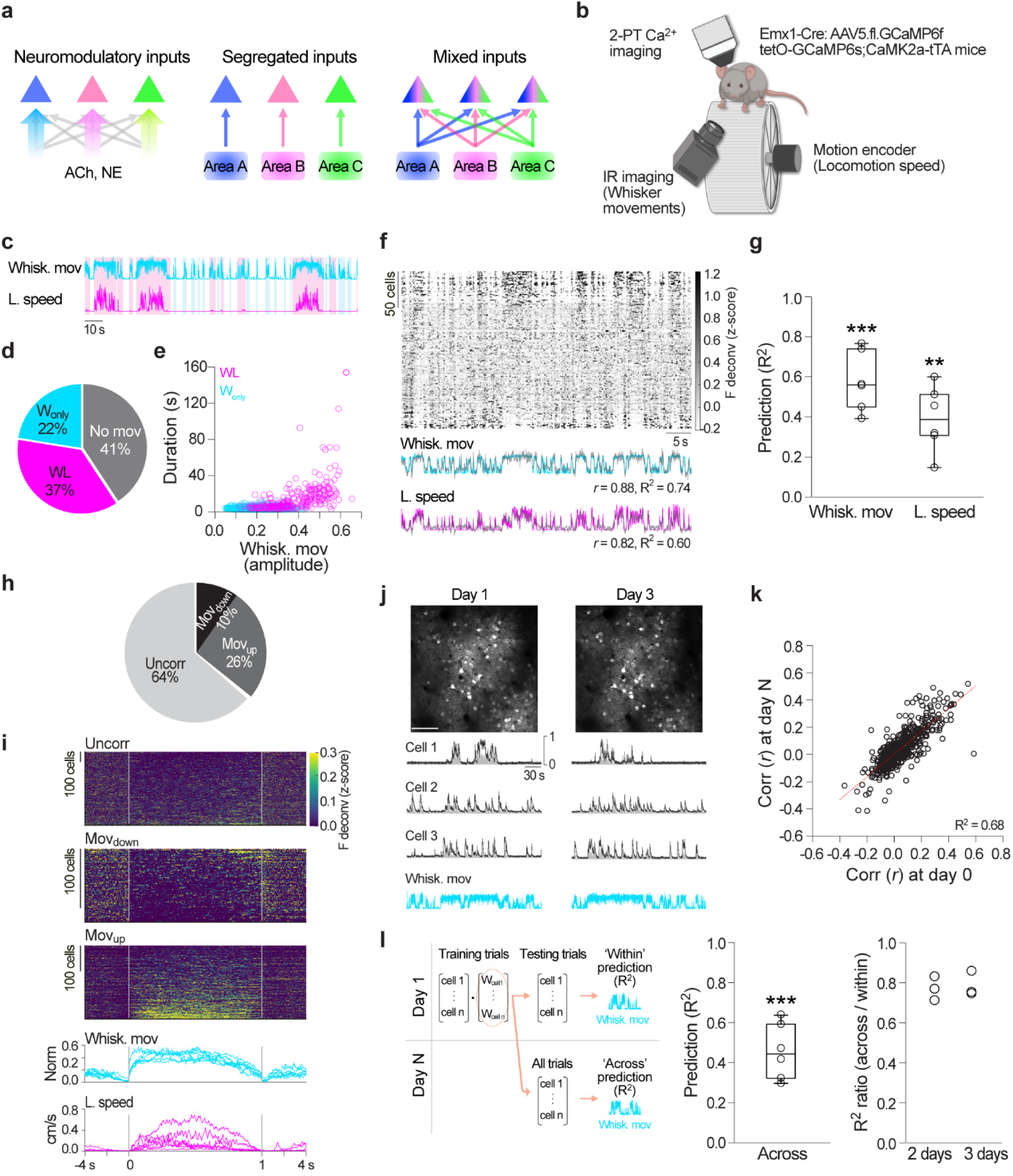
Heterogeneity and stability of neuronal activity patterns in relation to spontaneous movements. **a**, Hypothetical input connectivity models underlying the functional heterogeneity of cortical PNs. PNs with distinct activity patterns are represented by triangles of different colors. Left, heterogeneity is achieved through structured presynaptic neuromodulatory inputs. Middle, heterogeneity arises from a specific set of glutamatergic and/or GABAergic presynaptic inputs. Right, the functional properties of neurons are flexible, and their presynaptic networks are anatomically similar. **b**, Illustration of the experimental setup for 2-PT Ca^2+^ imaging of wS1 L2/3 PNs in head-fixed, awake mice. **c**, Example time course of extracted spontaneous movement variables, whisker movements (whisk. mov.) and locomotion (l.) speed. Spontaneous movement types: whisker movements without locomotion (W_only_, shaded, blue) and whisker movements with locomotion (WL, shaded, pink). Note that locomotion is always accompanied by whisker movements. **d**, Pie chart of time spent per spontaneous movement type (W_only_, WL) and not moving (no mov., *n* = 5 sessions, 5 mice). **e**, Relationship between whisker movement amplitude and duration for W_only_ (1268) and WL (321) events. **f**, Top, example raster plot of neuronal activity during spontaneous movements. Individual neurons (201) are sorted from top to bottom by decreasing weight on the first principal component. Bottom, prediction of either whisker movements or locomotion speed from population activity using two separate decoders. Correlation (corr., *r*) between predicted (gray) and actual whisker movement (blue, *P* < 0.0001) and locomotion speed traces (magenta, *P* < 0.0001), and variance explained (R^2^) by each model. **g**, Prediction of whisker movements and locomotion speed from neuronal activity. Variance explained within session (whisk. mov., *P* = 0.00011; l. speed, *P* = 0.0010; paired sample *t*-test R^2^ ≤ 0 vs. R^2^ > 0; *n* = 6 FOVs, 6 sessions, 5 mice). **h**, Pie chart of the relative proportions of movement-uncorrelated (uncorr., 715), Mov_down_ (123) and Mov_up_ (286) neurons. **i**, Peri-event time histogram (PETH) of movement-uncorrelated, Mov_down_, and Mov_up_ neurons for WL events that are time-normalized from onset to offset (vertical bars). **j**, Top, example imaging FOV at day 1 and day 3. Scale bar, 100 µm. Bottom, Activity of example Mov_up_ neurons and corresponding whisker movement traces at different days. Black, F; gray, F deconv.; both normalized to maximum. **k**, Scatter plot of correlation (*r*) between the activity of individual neurons and whisker movements at day 3 or day 4 vs. day 1 (*P* < 0.0001, regression; *n* = 682, 6 FOVs, 2 sessions per FOV, 5 mice). **l**, Left, Linear decoder to test the stability of population activity in relation to whisker movements across days. Middle, Variance explained across days (*P* < 0.0001, paired sample *t*-test across R^2^ ≤ 0 vs. R^2^ > 0). Right, Out-of-sample R^2^ ratio (R^2^_across_ / R^2^_within_) for different time intervals between sessions (day 1 vs. day 3 and day 1 vs. day 4).

## Results

### Heterogeneity and stability of spontaneous movement-dependent activity patterns

We first characterized the neural representations of spontaneous movements in the whisker primary somatosensory cortex (wS1) of mice. We used two-photon (2-PT) Ca^2+^ imaging to monitor the activity of pyramidal neurons (PNs) of right hemisphere wS1 layer (L) 2/3 (Fig. 1b). We induced the expression of the Ca^2+^ sensor GCaMP6 in PNs by either injecting a virus carrying GCaMP6f in the wS1 of Emx1-IRES-Cre mice or crossing the CaMK2a-tTA and tetO-GCaMP6s transgenic mouse lines. Mice were head-fixed on top of a wheel in darkness. A near-infrared camera was used to capture whisker movements (whisk. mov.) (Fig. 1b). A wheel speed encoder was used to measure locomotion (l.) speed (Fig. 1b). Mice were not instructed to move, nor were they rewarded. We observed two types of spontaneous movements throughout the recording session. One type consisted of short duration and small amplitude whisker movements without locomotion (W_only_). The other type consisted of longer and larger amplitude whisker movements accompanied by locomotion (WL) (Fig. 1c-e). To assess the relationship between neuronal activity and spontaneous movements, we trained two separate linear decoders to predict whisker movements or locomotion speed from population activity and evaluated their predictive performance. Population activity exhibited fluctuations coupled to the presence or absence of spontaneous movements and reliably predicted both whisker movements and locomotion speed in out-of-sample data (Fig. 1f-g). Since the fraction of variance explained by neuronal activity was higher for whisker movements than for locomotion speed (Fig. 1g), we based our population analysis on the whisker movement prediction^8^. Embedded in the population, individual neurons exhibited heterogenous patterns of activity in relation to spontaneous movements (Fig. 1f and Extended Data Fig. 1). Two subsets of neurons exhibited activity patterns time-locked to spontaneous movements. The first showed increased activity (Mov_up_, 26 ± 6.7% of all neurons) and the second, decreased activity during spontaneous movements (Mov_down_, 10 ± 6.3% of all neurons) (Fig. 1h-i). Most Mov_up_ neurons (75 ± 19% of all neurons) increased their activity during WL bouts and are hereafter referred to as movement-correlated neurons (corr.) (Fig. 1i and Extended Data Fig. 1). These results are consistent with the interpretation that wS1 is strongly engaged during locomotion^8^. All other neurons did not change their activity with respect to spontaneous movements (movement-uncorrelated neurons, uncorr.).

To understand how the spontaneous movement-correlated and sensory stimulus-responsive neuronal populations are represented in wS1, we recorded not only ongoing, but also sensory stimulus-evoked neuronal activity. Sensory stimulation consisted of a periodic deflection of the left whiskers, using a vertically oriented pole. We found that 10 ± 5.7% of all neurons responded positively to passive whisker deflection (Extended Data Fig. 2)^24,38^. A fraction of movement-uncorrelated (9.4%), Mov_up_ (14%), and Mov_down_ (7.9%) neurons responded to sensory stimulation (Extended Data Fig. 2), as observed previously in the visual cortex^11^ and wS1^8,24^. Together, these results show that a subset of wS1 PNs reliably encodes spontaneous movements and that the coding of spontaneous movements and sensory stimuli is independent given that some neurons have both properties.

We next asked whether the representation of spontaneous movements in wS1 PNs is a stable feature. While both stability and plasticity of sensory coding schemes^39^ and spontaneous ensemble activity^40^ in primary sensory cortices have been reported, the stability of spontaneous movement-dependent patterns of activity over the course of multiple days has not been directly tested. When tracking the same population of neurons over multiple days, we observed a highly stable correlation between neuronal activity patterns and spontaneous movements (Fig. 1j), both at single-neuron and population levels. The correlation (*r*) between the activity of individual neurons and whisker movements was highly reliable across days (Fig. 1k). To evaluate the stability of population activity in relation to spontaneous movements across days, we built a linear decoder using data from the first imaging session (day 1, within, Fig. 1l) and applied this decoder to out-of-sample data collected 2-3 days later (day N, across days). The day 1 model consistently predicted spontaneous movements across days (Fig. 1l), suggesting a stable representation in the population. Our results demonstrate that wS1 PNs reliably represent spontaneous movements over multiple days.

### Direct neuromodulatory inputs to spontaneous movement activity

We next investigated the source of the stable encoding of spontaneous movements by wS1 PNs. Neuromodulators, including neuropeptides, exert powerful effects over cortical function^34,41–43^. Acetylcholine (Ach) and norepinephrine (NE) terminal activity in sensory cortices, for example, is highly correlated to spontaneous movements^32,36,44–47^. These neuromodulators have been tightly linked to the locomotion-related gain modulation of sensory responses in PNs^31,48,49^. However, a direct effect of ACh and NE on the activity of large populations of L2/3 PNs during spontaneous movements remains to be established. To that end, we implanted a custom-made cranial window with an access port that allowed local application of ACh, NE, or glutamate receptor (R) blockers while imaging wS1 L2/3 PNs (Fig 2a). We blocked a different type of receptor or no receptor at all (in sham controls) per imaging session, and each session was performed on a different day. In a final session, tetrodotoxin (TTX), a Na^+^ channel blocker, was applied to evaluate the effectiveness of drug diffusion per FOV. TTX silenced nearly all neuronal activity over the entire FOV within 20 min (Extended Data Fig. 3). To further test whether ACh and NE R blockers reach all neurons in the FOV, we expressed a genetically engineered ACh (GRAB_ACh_) or NE (GRAB_NE_) sensor in wS1 L2/3 neurons, through viral vectors^45,47^. We observed increases in both ACh and NE R binding during spontaneous movements^32,36,44–47^. However, upon application of either ACh or NE R blockers, these increases were abolished throughout the FOV, demonstrating the effectiveness of the blocker application approach (Extended Data Fig. 4).

**Fig. 2:**
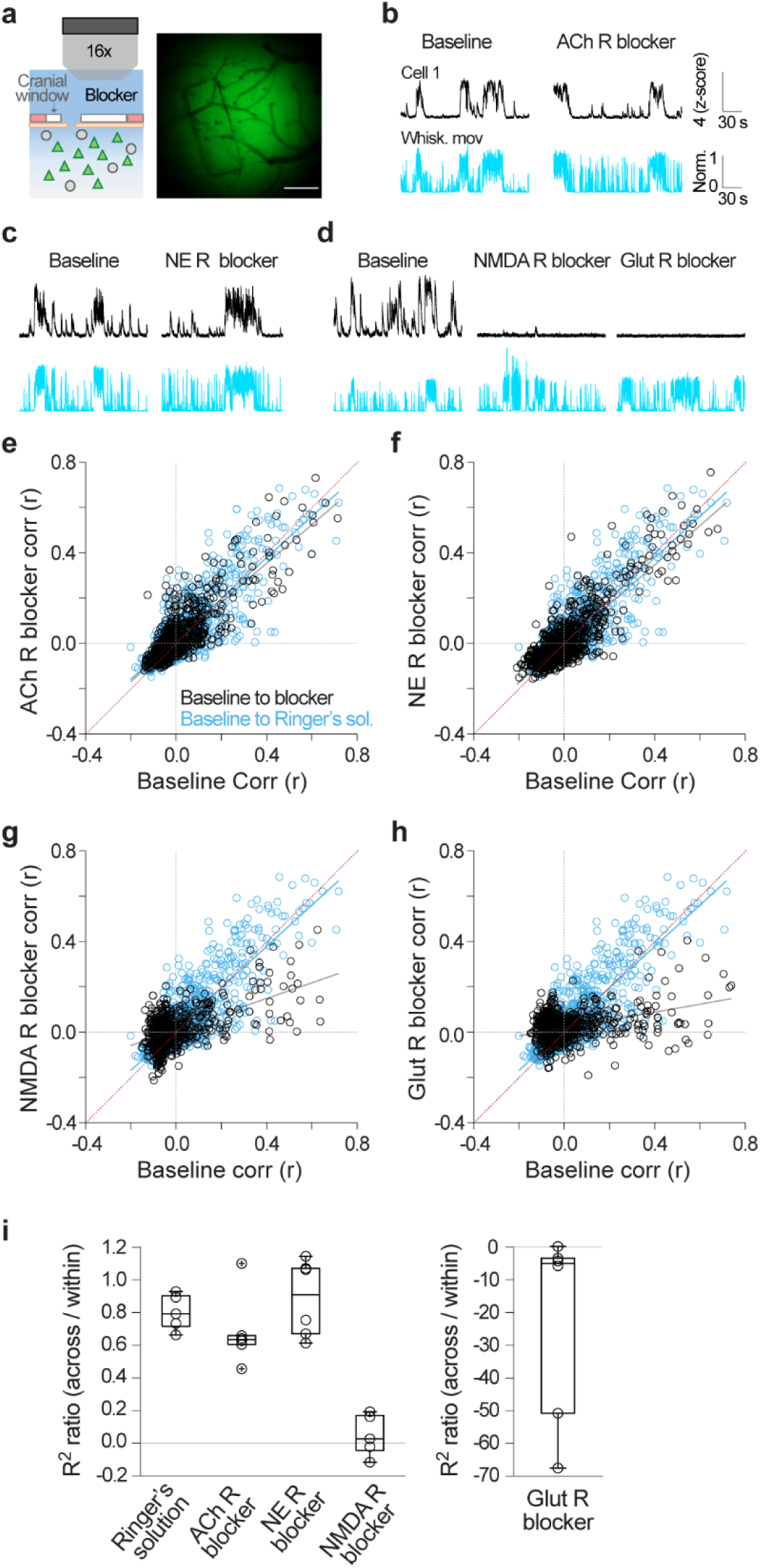
Limited role of direct neuromodulatory inputs for spontaneous movement-dependent neuronal activity in wS1. **a**, Left, Experimental approach for local delivery of R blockers during 2-PT Ca^2+^ imaging using a custom cranial window. Imaging was performed without (baseline) or with R blockers in the Ringer’s solution. In sham sessions, following baseline imaging, the Ringer’s solution was replaced by Ringer’s solution without R blockers. Triangles represent PNs. Circles represent GABAergic neurons. Right, Example cranial window with a laser-cut opening. Scale bar, 0.3 mm. **b-d**, Activity of an example Mov_up_ neuron (F, black), before and after pharmacological blockade of ACh (atropine and mecamylamine, **b**), NE (prazosin and propranolol, **c**), NMDA (D-AP5, **d**), or NMDA and AMPA (Glut, D-AP5 and DNQX, **d**) R; ACh, NE and NMDA/Glut R were tested on different days. Blue traces, whisker movements. **e-h**, Scatter plots of the correlation (*r*) of individual neurons with whisker movements during ACh (*n* = 1300, 6 sessions, 6 mice, **e**), NE (*n* = 1483, 6 sessions, 6 mice, **f**), NMDA (*n* = 720, 5 sessions, 4 mice, **g**), or Glut (*n* = 956, 6 sessions, 6 mice, **h**) R blockade vs. baseline, respectively; sham sessions (baseline vs. Ringer’s only) are included in e-h (cyan, *n* = 1598, 5 sessions, 3 mice) (null hypothesis, equal slopes: ACh R vs. sham, *P* = 0.033; NE R vs. sham, *P* = 0.13; NMDA R vs. sham, *P* < 0.0001; Glut R vs. sham, *P* < 0.0001; multiple comparisons). **i**, Prediction of whisker movements from population activity. Out-of-sample R^2^ ratio (R_across_ / R_within_) (ACh R vs. sham, *P* = 0.33; NE R vs. sham, *P* > 0.05; NMDA R vs. sham, *P* = 0.032; Glut R vs. sham, *P* = 0.017; NMDA R vs. Glut R, *P* < 0.0001; Kruskal-Wallis test (*P* < 0.001) followed by Wilcoxon rank-sum tests).

The temporal correlation between the activity of individual neurons and spontaneous movements was largely preserved during ACh or NE R blockade (Fig. 2b-c, e-f), suggesting that individual neurons maintain their activity profile during spontaneous movements. However, application of an NMDA R blocker diminished the correlation between individual neuron activity and spontaneous movements, revealing a strong NMDA-component of spontaneous movement-dependent activity (Fig. 2d, g). The combined action of NMDA and AMPA (Glut) R blockers further uncovered the contribution of glutamatergic signaling through AMPA R for ongoing activity (Fig. 2d, h and Extended Data Fig. 5). While ACh and NE R blockers did not disrupt the correlation of PNs with spontaneous movements, both blockers modulated the magnitude of movement-dependent activity in a fraction of movement-correlated neurons. The effects of the blockers were higher on evoked activity in stimulus-responsive neurons (Extended Data Fig. 5).

We examined the effect of neuromodulatory and glutamatergic inputs on population activity. For each recording session, we built a linear decoder using data recorded prior to the application of the receptor blocker(s) or Ringer’s reapplication in sham controls (within), then tested this decoder using the out-of-sample data acquired after blocker application (across conditions). The model consistently predicted spontaneous movements from population activity across conditions when either ACh or NE R blockers were applied, indicating that the structure of movement-dependent population activity is largely maintained during neuromodulator receptor blockade (Ringer’s sham controls, R^2^ = 0.54 ± 0.085; ACh R, R^2^ = 0.37 ± 0.084; NE R, R^2^ = 0.52 ± 0.15; *P* < 0.0001 for all conditions, paired sample *t*-test across R^2^ ≤ 0 vs. R^2^ > 0). By contrast, the decoder failed to predict spontaneous movements from population activity when either NMDA or Glut R blockers were tested (NMDA R, R^2^ = 0.025 ± 0.084, *P* = 0.26; Glut R, R^2^ = -11.32 ± 15, *P* = 0.95). The out-of-sample R^2^ ratio (R^2^_across_ / R^2^_within_) of ACh or NE R blockade did not differ from that of the sham controls, but it was significantly lower for NMDA and Glut R blockade (Fig. 2i). Local application of the different blockers did not affect spontaneous movement behavior (Extended Data Fig. 6). These results provide evidence for a limited role of direct neuromodulatory (ACh and NE) inputs in generating highly stable patterns of spontaneous movement-dependent activity in wS1.

### Single-cell based monosynaptic retrograde tracing

Based on the highly stable, glutamate receptor-sensitive activity during spontaneous movements, we hypothesized that a distinct anatomical organization of presynaptic networks may constrain the spontaneous movement-dependent activity profiles of PNs. To reveal the presynaptic network of functionally identified neurons at the single-cell level, we adapted a single-neuron based monosynaptic retrograde tracing approach^16,19,50–53^ (Fig. 3a). We first imaged wS1 L2/3 PNs and selected a target neuron based on its activity profile during spontaneous movements (example movement-correlated neuron, Fig. 3a, b). We then performed 2-PT guided electroporation of the target neuron with DNA encoding for TVA (rabies virus (RV) receptor), G (RV spike protein), and mCherry (for validation of transfection) (Fig. 3a and Extended Data Fig. 7). Following electroporation, we injected a RV variant carrying a red fluorescent protein (RFP, mCherry *n* = 2 brains, tdTomato (tdTom), *n* = 20 brains) close to the electroporated neuron. In line with previous studies^16,19^, at day 4-5 we observed the emergence of RFP^+^ presynaptic neurons in wS1. We detected the RFP^+^ neurons exclusively in brains where the electroporated neuron survived for several days and expressed mCherry (postsynaptic neuron), confirming the specificity of our approach (Extended Data Fig. 8). To reveal presynaptic neurons, animals were perfused at day 11 (+/-1.5 days), and whole brains were analyzed histologically. Presynaptic neurons were manually annotated according to anatomical area and identified as glutamatergic or GABAergic based on the immunostaining with GABA (see Methods, Whole-brain reconstruction, annotation, and registration). Brains were registered to the Allen Mouse Common Coordinate Framework, and annotations were validated (Fig. 3c).

**Fig. 3:**
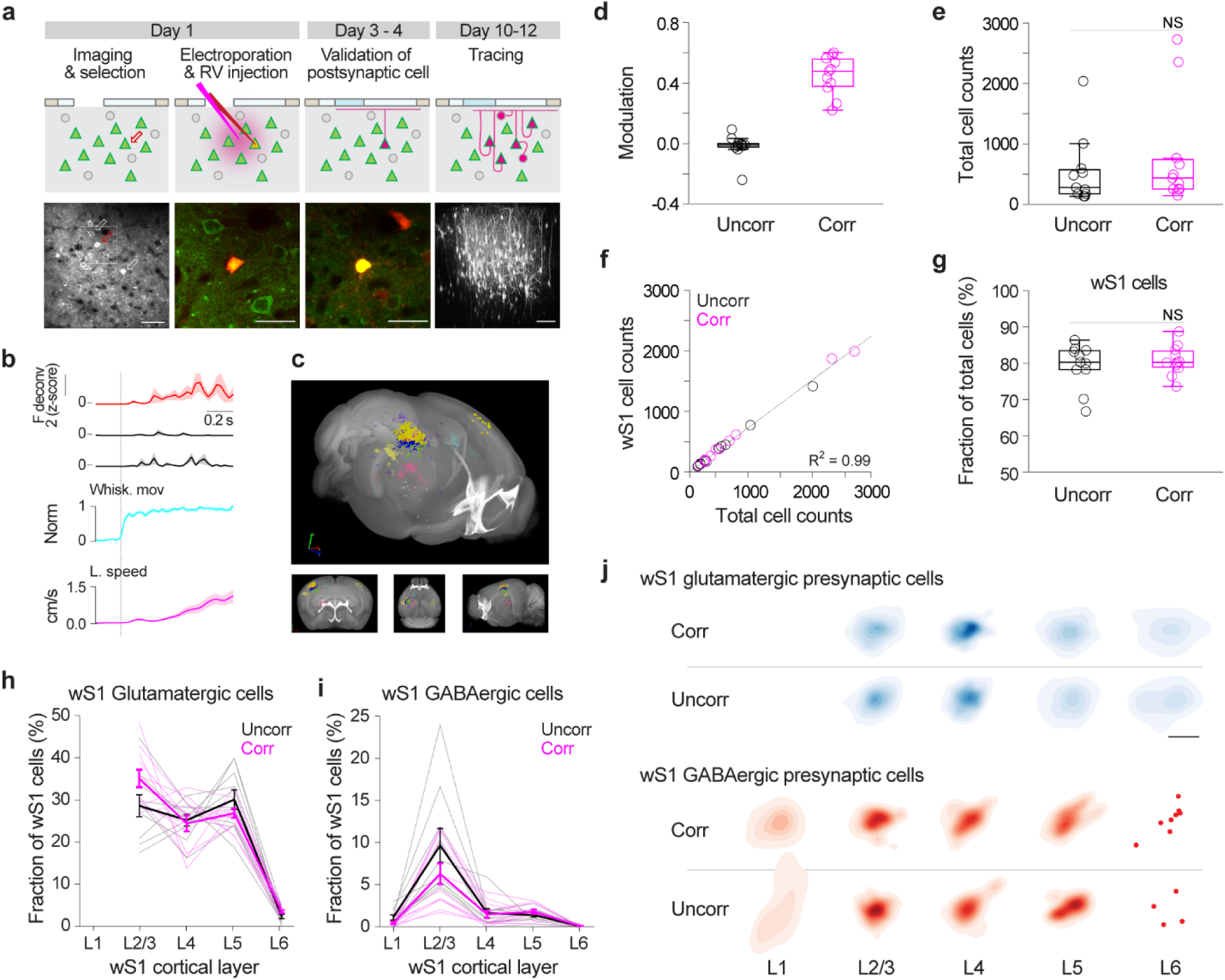
Functionally distinct neurons have anatomically similar local presynaptic networks. **a**, Experimental procedure for tracing monosynaptic inputs to a single, functionally identified PN. Day 1: Left, Imaging and selection of a target neuron (red arrow) based on its activity profile with respect to spontaneous movements. Example FOV. Scale bar, 50 µm. Right, 2-PT guided electroporation of the target neuron and subsequent injection of RV-RFP. Image of the target neuron immediately after electroporation (GCaMP6s^+^-Alexa594^+^). Scale bar, 25 µm. Day 3-4: Evaluation of survival and transfection. Image of the target neuron expressing mCherry (GCaMP6s^+^-mCherry^+^). Scale bar, 25 µm. Day 10-12: Emergence of the local presynaptic network (RFP^+^). Three-dimensional projection of z-stacks (814 x 814 x 785 µm) of wS1 presynaptic neurons (RFP^+^). Scale bar, 100 µm. **b**, Activity (F deconv.) of the example target neuron (red) and two surrounding neurons (black, indicated by white arrows in a) aligned to the onset of WL events (mean ± s.e.m.). **c**, Brain-wide presynaptic network of the example postsynaptic neuron (shown in **a** and **b**) in the Allen Mouse Common Coordinate Framework. Presynaptic neurons were anatomically parsed. **d**, Mean modulation of all postsynaptic neurons during WL events (11 movement-uncorrelated, uncorr., neurons from 11 mice, and 11 movement-correlated, corr., neurons from 11 mice, 1 neuron per mouse). **e**, Total number of presynaptic neurons per brain for each postsynaptic neuron group (*P* = 0.39, randomization test). **f**, Scatter plot of wS1 vs. total presynaptic neurons (*P* < 0.0001, regression). **g**, Fraction of presynaptic neurons in wS1 (*P* = 0.52, randomization test). **h**, Distribution of wS1 glutamatergic presynaptic neurons of movement-uncorrelated (*n* = 10) and movement-correlated (*n* = 11) postsynaptic neurons across cortical layers (layer vs. layer, L6 vs. all other layers, *P* < 0.01, Kruskal-Wallis test (*P* < 0.0001) followed by Dunn’s test; uncorr. vs. corr., *P*_L2/3_ = 0.065, *P*_L4_ = 0.74, *P*_L5_ = 0.20, *P*_L6_ = 0.12, randomization tests). Thin lines, individual brains; thick lines, mean ± s.e.m.. **i**, Distribution of wS1 GABAergic presynaptic neurons (layer vs. layer, L2/3 vs. all other layers, *P* < 0.01, Kruskal-Wallis test (*P* < 0.0001) followed by Dunn’s test; GABAergic, uncorr. vs. corr., *P*_L2/3_ = 0.16; *P*_L4_ = 0.79; *P*_L5_ = 0.28, randomization tests). **j**, Horizonal projections of the weighted distributions of wS1 glutamatergic or GABAergic presynaptic neurons per cortical layer (glutamatergic, uncorr. vs. corr., *P*_L2/3_ = 0.73, *P*_L4_ = 0.39, *P*_L5_ = 0.066, *P*_L6_ = 0.26; GABAergic, uncorr. vs. corr.*, P*_L2/3_ = 0.86; *P*_L4_ = 0.84; *P*_L5_ = 0.31; L2/3 glutamatergic vs. GABAergic average pairwise distance between neurons, *P* = 2.5 x 10^-09^, *t*-test). Scale bar, 500 µm.

We performed single-cell based retrograde tracing of two functionally distinct subsets of neurons: movement-uncorrelated (*n* = 11, 11 mice) and movement-correlated (*n* = 11, 11 mice) neurons (Fig. 3b, d). To refine the functional specificity and eliminate the confounding factor of somatosensory input in both groups, we targeted neurons that did not respond to sensory stimulation. Spontaneous movements, the proportion of spontaneous movement-dependent neuronal subsets, and cortical depth of the postsynaptic (target) neurons were similar between the two groups (Extended Data Fig. 9).

To analyze the brain-wide arrangement of presynaptic connectivity, we compared the fraction of presynaptic cells from each brain area between the two groups. Variability in the total number of presynaptic neurons across brains is expected due to differences in the survival time of each postsynaptic neuron^54^. Yet, the distribution of the total number of presynaptic neurons per brain was comparable for the movement-uncorrelated (range = 137-2038) and movement-correlated groups (range = 148-2727) (Fig. 3e). We observed a strong linear association between the number of local vs. total presynaptic neurons per brain in both groups (Fig. 3f and Extended Data Fig. 10). Based on the linear association, we concluded that comparing fractional presynaptic inputs onto postsynaptic neurons of the two groups is appropriate.

### Local (wS1) presynaptic networks

We first investigated whether spontaneous movement-correlated neurons receive inputs from distinctive local, wS1 presynaptic networks. Presynaptic neurons located in wS1 constituted the largest fraction of brain-wide inputs to each postsynaptic neuron: 79 ± 6.0% for the movement-uncorrelated group and 81 ± 6.0% for the movement-correlated group (Fig. 3g and Extended Data Fig. 10). Of all wS1 presynaptic neurons, the majority were glutamatergic: 86 ± 7.4% for the movement-uncorrelated and 90 ± 5.5% for the movement-correlated group (Extended Data Fig. 10). Glutamatergic presynaptic neurons were, on average, broadly distributed across L2/3 to L5 but significantly less predominant in L6 for both groups (Fig. 3h). Individual glutamatergic presynaptic networks were often characterized by a smaller fraction of L4 compared to L2/3 or L5 neurons^55,56^. The predominance of this laminar profile was comparable between the groups (*P* = 0.95, randomization test with Chow test). Additionally, the fraction of glutamatergic inputs from each layer was similar between the groups. In contrast to the broad vertical distribution of glutamatergic inputs, GABAergic presynaptic neurons were mostly limited to L2/3 (Fig. 3i)^19^. Yet, the fraction of GABAergic inputs per layer did not differ between the two groups. These results demonstrate that the functionally distinct neurons have a similar proportion of local presynaptic neurons across layers.

Next, we explored the spatial distributions of wS1 presynaptic networks. The three-dimensional and two-dimensional (layer-by-layer horizontal flat projections) spatial distributions of wS1 glutamatergic and GABAergic presynaptic networks were not significantly different between the groups (Fig. 3j). To further characterize the horizontal spread of wS1 glutamatergic presynaptic networks across layers, we performed gaussian density estimation on the layer projections (Extended Data Fig. 11). glutamatergic presynaptic neurons in L4 were restricted to a smaller cortical span (cell pairwise distance mean, ∼309 µm) than in L2/3 or L5 (mean, ∼417 and 505 µm, respectively), consistent with the notion that L2/3 neurons receive inputs from mainly one barrel^57,58^. We observed this feature in all 22 presynaptic networks of both groups. GABAergic presynaptic neurons were more spatially confined than glutamatergic presynaptic neurons^19^ (cell pairwise distance mean for L2/3, 272 ± 71 µm) (Fig. 3j). In summary, the glutamatergic and GABAergic presynaptic networks of single L2/3 PNs in wS1 exhibit anatomical features found in population studies^55–58^. Yet, despite their markedly different activity patterns, functionally distinct sets of neurons (movement-uncorrelated and movement-correlated) have a similar pattern of local glutamatergic and GABAergic inputs, not only in terms of number but also spatial distribution across all cortical layers.

### Long-range presynaptic ensembles

We next investigated whether movement-correlated neurons receive a distinctive set of brain-wide, long-range inputs (Fig. 1a). Recent studies provided insights into the organization of local inputs onto individual PNs in primary visual cortex in the context of visual stimuli^16,19^. Yet, how the heterogenous activity patterns of L2/3 neurons relate to long-range inputs has not been explored. Analysis of the single-cell based, whole-brain presynaptic networks revealed the distribution of presynaptic neurons in multiple cortical and subcortical brain areas that are known to project to wS1^55,56,59^: secondary somatosensory cortex (S2), primary and secondary motor cortices (M1/2), thalamus, other sensory cortical areas (SenCtx, including auditory visual cortices), contralateral wS1 (cwS1), perirhinal cortex (PrhCtx), and basal forebrain (BF) (Fig. 4a). Brain areas containing, on average, less than 0.5% of total presynaptic cells were grouped into ‘Others’. All long-range presynaptic neurons were glutamatergic. Most input areas were ipsilateral, with a few exceptions such as the PrhCtx. Irrespective of its activity pattern, each postsynaptic neuron from both the movement-uncorrelated and movement-correlated groups received inputs from, on average, 6.5 of the 7 major brain areas known to project to wS1 (Fig. 4a, b). These results suggest that a high degree of integration or multiplexing can occur at the single-cell level. While all postsynaptic cells from both groups received direct inputs from a wide range wS1-projecting brain areas, we found that, surprisingly, movement-correlated neurons receive a lower fraction of inputs from M1/2 than movement-uncorrelated neurons (Fig. 4a, c). In contrast, movement-correlated neurons received a significantly higher fraction of inputs from thalamus relative to movement-uncorrelated neurons (Fig. 4a, d). Thalamic presynaptic neurons were almost equally distributed in the first order thalamic relay nucleus, ventral posteromedial nucleus (VPm), and the higher order thalamic nucleus, posteromedial nucleus (POm) for both groups (Extended Data Fig. 12). Similarly, the fraction and spatial distribution of M1 and M2 cells in motor cortical presynaptic networks were comparable between the groups (Extended Data Fig. 12). Furthermore, the modulation of the activity of postsynaptic neurons across spontaneous movements (i.e., change in activity during movement relative to baseline, prior to movement onset, see Methods, Modulation of neuronal activity) was negatively correlated with the fraction of presynaptic neurons found in M1/2 (Fig. 4e) and positively correlated with the fraction of presynaptic neurons found in thalamus (Fig. 4f). Within each long-range input area, the spatial distribution of presynaptic neurons was similar for the two groups (Extended Data Fig. 13), suggesting that the functionally distinct postsynaptic neurons receive inputs from spatially intermingled long-range neurons. Overall, these results demonstrate that, despite the high degree of convergence of inputs from multiple brain areas to single neurons in wS1, movement-correlated wS1 PNs show characteristic brain-wide presynaptic inputs, with a relatively larger fraction of thalamic, but lower fraction of motor cortical inputs.

**Fig. 4:**
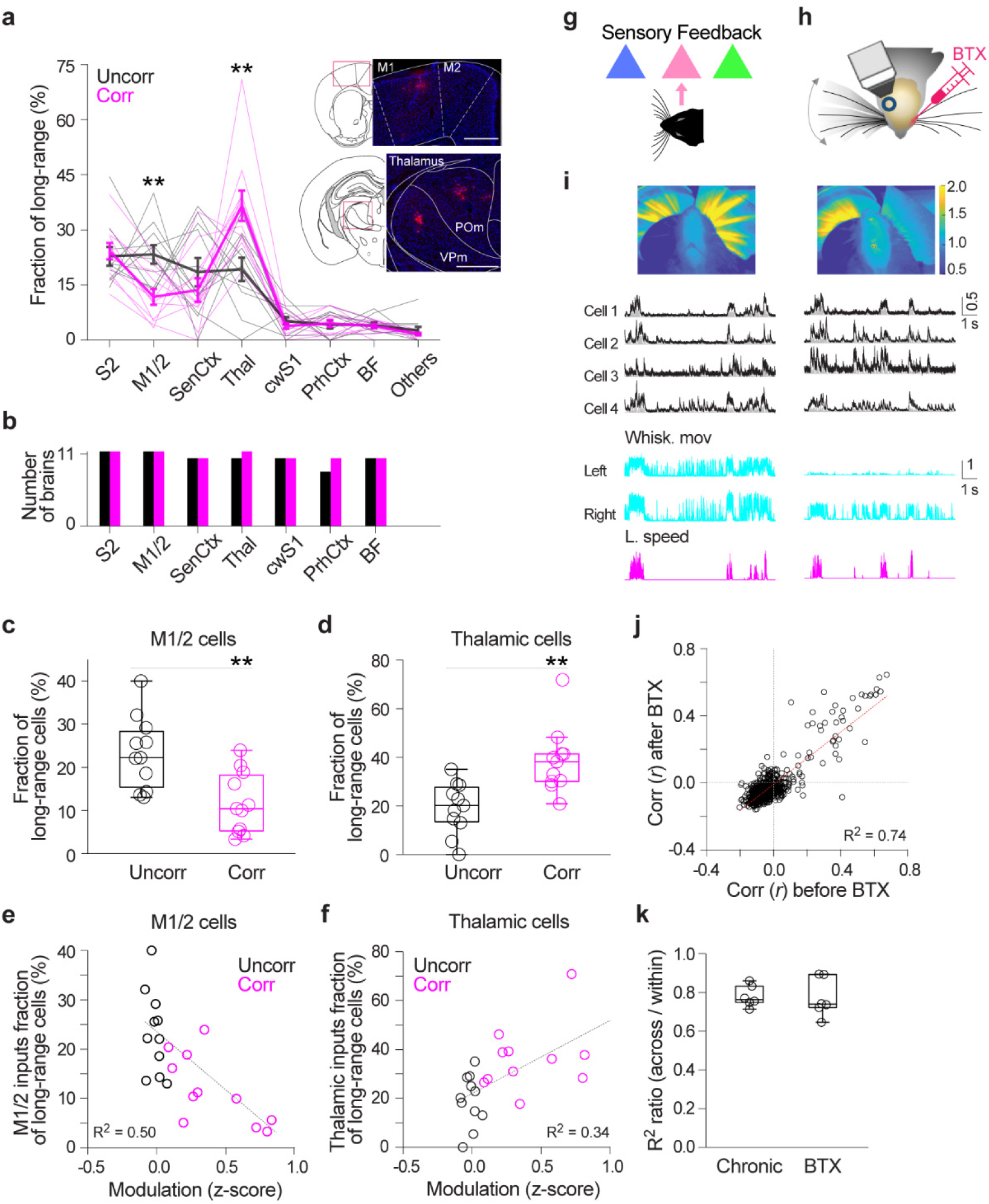
Functionally distinct neurons receive characteristic long-range inputs. **a**, Distribution of the presynaptic network of each movement-uncorrelated (*n* = 11) or movement-correlated postsynaptic neuron (*n* = 11) across multiple brain areas (M1/2, *P* = 0.0030; thalamus, *P* = 0.0030; all other areas, *P* > 0.05, randomization tests). Thin lines, individual brains; thick lines, mean ± s.e.m.. Inset, example epifluorescence images of coronal brain slices denoting the presence of presynaptic neurons (RFP^+^) in M1/2 and thalamus (predominantly in VPm and POm) (from example presynaptic network shown in Fig. 3a, d). Scale bar, 500 µm. **b**, Number of brains containing presynaptic neurons in each of the listed brain areas. Black, uncorr. group, magenta, corr. group. Note the high degree of convergence of inputs from diverse areas at the single-cell level. **c-d**, Motor cortical (**c**) and thalamic (**d**) presynaptic neurons. **e-f**, Motor cortical (**e**) and thalamic (**f**) input fraction as function of the average modulation of the postsynaptic neurons across spontaneous movements (W_only_ + WL) (M1/2, *P* = 0.0024; thalamus, *P* = 0.0045; regression). **g**, Sensory feedback from whisker movements as a hypothetical source of movement-dependent activity in a subset of PNs in wS1. **h**, Schematic of imaging over the right hemisphere wS1 before (left) and 1-2 days (right) after BTX injection in the left mystacial pad. **i**, Top, Example images of videorecorded whisker movements (mean of the absolute difference between consecutive frames over ∼2 min), before and after BTX injection. Warmer colors reflect a higher motion. The BTX injection effectively paralyzed the left mystacial pad (*n* = 5 mice). Bottom, Example neuronal activity with corresponding whisker movement and locomotion speed traces before and after BTX injection. Black, F, and gray, F deconv., both normalized to maximum. **j**, Correlation (r) of the activity of individual neurons with whisker movements before vs. after BTX injection (*P* < 0.0001, regression; *n* = 1044, 6 FOVs, 2 sessions per FOV, 5 mice). **k**, Prediction of whisker movements from population activity. Out-of-sample R^2^ ratio (R^2^_across_ / R^2^_within_) for chronic (Fig. 1) vs. BTX experiments (*P* = 0.70, Wilcoxon rank-sum test).

Given that movement-correlated neurons receive more abundant inputs from the thalamus, we next asked whether the spontaneous movement-dependent activity in these neurons is caused by sensory feedback from voluntary whisker movements (Fig. 4g). We imaged right hemisphere wS1 L2/3 PNs before and after paralysis of the left facial muscles by injection of botulinum toxin (BTX) in the left mystacial pad (Fig. 4h). The same population of PNs was reimaged 2-3 days after BTX injection. This procedure was minimally invasive but effectively abolished left whisker movements (Fig. 4i) without interrupting right whisker movements and locomotion (Extended Data Fig. 14). Since left and right whisker movements were highly correlated (*r* left vs. right = 0.959 ± 0.0165, *n* = 4 sessions, 4 mice), we used the right whisker movements to detect changes in movement-dependent activity after paralysis (Fig. 4i). We observed that the correlation between the activity of individual neurons and spontaneous movements was highly preserved even after paralysis (Fig. 4j). Beyond correlation, we found that the modulation of individual neurons during spontaneous movements was similar before and after paralysis (Extended Data Fig. 14). To examine changes in the relationship between population activity and spontaneous movements, we built a linear decoder using data collected prior to paralysis (within) and evaluated it on out-of-sample data recorded after paralysis (across days). Model predictions were highly reliable across days (R^2^ = 0.44 ± 0.092, *P* < 0.0001, paired sample *t*-test test across R^2^ ≤ 0 vs. R^2^ > 0). In addition, the out-of-sample R^2^ ratio (R^2^_across_ / R^2^_within_) was similar when recordings were performed across days without (Fig. 1, chronic) and with unilateral whisker paralysis (Fig. 4k). To further eliminate a potential contribution of sensory feedback from the ipsilateral mystacial pad, we monitored neuronal activity in wS1 following bilateral whisker trimming, as well as combined bilateral whisker trimming and mystacial pad paralysis. We found that spontaneous movement-dependent activity is largely preserved even after combined bilateral whisker trimming and mystacial pad paralysis (Extended Data Fig. 15). These results revealed that the individual neuron and structure of population activity in relation to spontaneous movements are stable after facial paralysis and whisker trimming, suggesting that sensory feedback evoked by whisker movements does not play an instrumental role in driving spontaneous movement-dependent activity in wS1.

### Suppression of long-range thalamic inputs

Finally, we asked if the anatomical signature of predominant thalamic inputs to movement-correlated neurons contributes to their activity during spontaneous movements. We imaged PNs while simultaneously suppressing thalamic terminals in wS1 L2/3 (Fig 5a). To that end, we expressed the red-shifted inhibitory opsin archaerhodopsin ArchT, coupled to tdTomato, in the thalamus (VPm and POm) or M1/2 of tetO-GCaMP6s;CaMK2a-tTA mice (Fig. 5a-b). Mice that did not express ArchT served as controls. Light pulses (1-1.5 s) were randomly provided during the recording session. We observed that light pulses elicited a relatively brief (∼0.5 s) whisker movement in both control and ArchT-expressing mice (Extended Data Fig. 16). This whisker movement was independent of the ongoing movements of the animal and could potentially obfuscate neuronal recordings. To isolate changes in neuronal activity caused by thalamic terminal suppression from those caused by this light-evoked brief whisker movement, we restricted our single neuron analysis to the later light pulse window (0.5-1.0/1.5 s) during which the level of ongoing movements was similar to that of baseline (before the light pulse) (Extended Data Fig. 16). In ArchT-expressing mice, optogenetic suppression of thalamic terminals significantly altered the activity levels in a fraction of movement-correlated neurons (14%), producing a net decrease in their activity. This decrease was significantly larger than changes observed in movement-uncorrelated neurons (Fig. 5c-h). To further test the contribution from VPm and POm inputs, we expressed ArchT exclusively in either VPm or POm (Extended Data Fig. 17). Independent optogenetic suppression of VPm or POm terminals revealed that VPm predominantly contributes to the decreased activity of movement-correlated neurons. However, suppression of VPm terminals appears to produce a less robust effect than combined suppression of VPm and POm terminals. In M1/2 ArchT-expressing mice and in control mice, while we also detected changes in the activity of a fraction of movement-correlated neurons (7.6% and 10%, respectively), the net effect did not differ from that of movement-uncorrelated neurons. To address the optogenetic effect in a more comprehensive manner, we built a linear decoder using light-off periods and tested this model on data acquired during light presentation. We found a consistently larger decrease in R^2^ values for thalamic ArchT-expressing mice than for M1/2 ArchT-expressing or control mice (Fig. 5i-j). In some experiments, the across R^2^ value was close to zero, demonstrating that inhibition of thalamic inputs critically changed the relationship between population activity and spontaneous movements. These results provide evidence for the contribution of sensory feedback-independent, thalamic inputs, in particular from VPm, to spontaneous movement-dependent activity in cortical neurons.

**Fig. 5:**
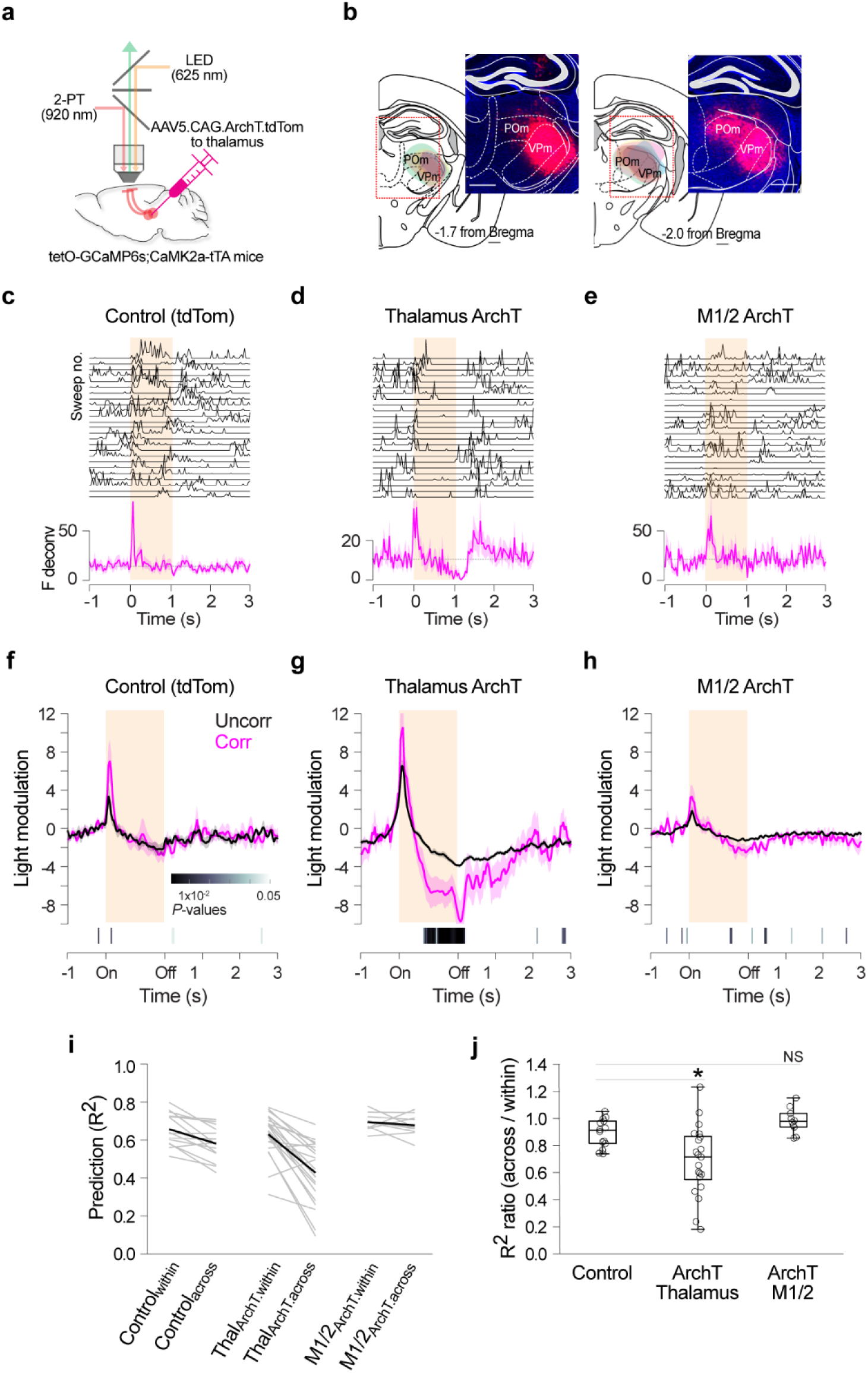
Optogenetic suppression of thalamic and motor cortical inputs on spontaneous movement-dependent activity. **a**, Schematic of the experimental approach for imaging of L2/3 PNs and simultaneous optogenetic suppression of thalamic and motor cortical axon terminals (ArchT^+^-tdTom^+^) in wS1. **b**, Example composite epifluorescence images of coronal brain sections showing tdTom^+^ cell bodies in VPm (left, AP -1.7 mm from Bregma), and VPm and POm (right, AP -2.0 mm from Bregma) in a thalamus ArchT-expressing mouse. ArchT-tdTom^+^ areas in thalamus are overlayed on the corresponding mouse atlas images; each color of shading represents an individual mouse (*n* = 7). Scale bars, 0.5 mm. **c-e**, Effect of light pulses on the activity of movement-correlated neurons. Example neurons from a control mouse (ArchT^-^-tdTom^+^, **c**), from a thalamus ArchT-expressing mouse (**d**), and from a M1/2 ArchT-expressing mouse (**e**). Top, responses to individual light pulses (F deconv.). Bottom, average PETH of baseline-subtracted activity. Orange shaded area indicates light pulse duration (time-normalized). **f-h**, Baseline-subtracted mean activity (light modulation) of movement-uncorrelated (black) and movement-correlated (magenta) neurons significantly affected by light pulses. *P* values, comparison of activity of movement-uncorrelated and movement-correlated neurons (Wilcoxon rank-sum test). **f**, Control mice (affected neurons, *n* = 30 of 184 uncorr. and *n* = 16 of 157 corr., 9 mice). **g,** Thalamus ArchT-expressing mice (affected neurons, *n* = 146 of 645 uncorr. and *n* = 25 of 182 corr., 7 mice). **h,** M1/2 ArchT-expressing mice (affected neurons, *n* = 39 of 271 uncorr. and *n* = 13 of corr., 3 mice). **i**, Prediction of whisker movements by population activity using a model trained on light-off data and evaluated on light-off (within) and light-on (across) data (explained variance). **j**, Out-of-sample R^2^ ratio (R^2^_across_/ R^2^_within_, thalamus ArchT vs. control, *P* = 0.012; M1/2 ArchT vs. control, *P* = 0.10; Kruskal-Wallis (*P* = 0.0005) followed by Wilcoxon rank-sum tests).

## Discussion

Here, we investigated anatomical wiring rules for the functional heterogeneity of cortical neurons. Our results provide insights into the functional and anatomical organization of cortical PNs in the context of behavioral state. We first demonstrated that the representation of spontaneous movements in wS1 PNs is stable over multiple days, both at the single-cell and population levels. What are the mechanisms underlying this stable, heterogenous representation? Based on the modest role of neuromodulatory inputs in wS1 and the decisive effect of glutamatergic inputs in driving this spontaneous movement-dependent activity, we investigated the anatomical architecture of brain-wide presynaptic networks of two subsets of wS1 L2/3 PNs (movement-uncorrelated & movement-correlated neurons). We found that individual PNs in superficial layers, regardless of their functional properties, receive highly converging inputs from most wS1-projecting brain areas, suggesting that individual cortical neurons have direct access to a broad range of information from diverse brain regions. Yet, functionally distinct cortical neurons showed anatomical biases in the proportions of specific long-range presynaptic inputs (Extended Data Fig. 18), suggesting that information from a given presynaptic area may be selectively amplified or reduced through the number of inputs.

Movement-correlated neurons received a lower fraction of motor cortical inputs and a higher fraction of thalamic inputs relative to movement-uncorrelated neurons. Moreover, we found a negative correlation between the modulation of neuronal activity by spontaneous movements and the fraction of motor cortical presynaptic cells and, conversely, a positive correlation between modulation of neuronal activity and fraction of thalamic presynaptic cells. Not only was the fraction of motor inputs onto movement-correlated neurons lower, but also optogenetic suppression of motor cortical axon terminals in wS1 had no effect on L2/3 PNs spontaneous movement-dependent activity. By contrast, in addition to an increased fraction of thalamic inputs converging onto movement-correlated neurons, we found that thalamic inputs directly contribute to the activity of PNs during spontaneous movements. A role for thalamic nuclei in driving behavioral-state dependent cortical activity is consistent with several earlier observations. First, robust manipulations of thalamic activity profoundly alter cortical state^60,61^. Second, thalamic nuclei activity correlates positively with spontaneous movements^10,11,29,61–65^, while the activity of neurons in cortical areas can fluctuate positively and negatively with spontaneous movements^11^. Third, thalamic inputs can drive L2/3 PNs more efficiently than cortical inputs^66^. However, that individual PNs encoding spontaneous movements have an enhanced thalamic input fraction could not have been predicted from previous studies. While movement-correlated PNs receive almost equally abundant inputs from VPm and POm, our optogenetic study demonstrates that the VPm nucleus is primarily responsible for driving the movement-dependent activity in wS1 L2/3. Eliminating whisker movements through muscle paralysis and sensory responses through whisker trimming did not disrupt spontaneous movement-dependent activity in wS1, strongly supporting the notion that this activity is unlikely to be a direct consequence of sensory feedback^60,65,67^. This raises an interesting issue: what do VPm neurons relay to the primary somatosensory cortex beyond simple sensory transmission, and how is the transmission of sensory and non-sensory information achieved? Sensory thalamic neurons, as cortical neurons, are also active in the absence of sensory stimuli and feedback^60^, suggesting that they can encode non-sensory information. Anatomical analysis supports the heterogeneous innervation patterns of individual VPm neurons across different cortical layers^68,69^. Future studies will be needed to address the potential functional heterogeneity in VPm neurons^64^.

The release of ACh and NE in cortex is closely linked to spontaneous movements, as also confirmed in our GRAB sensor experiments^32,36,44–47^. Our results shows that these inputs may influence the activity levels, especially with respect to sensory responses, consistent with the gain modulation of sensory responses during movement by neuromodulation^31,48,49^. However, our neuromodulatory blockade results suggest that the direct neuromodulatory inputs to the cortex are not the main drivers for the neuronal activity in relation to spontaneous movements. Instead, the subset of neurons that reliably tracked the behavioral state appeared to be driven by long-range glutamatergic inputs, particularly from the thalamus. Neuromodulators profoundly alter the thalamic activity mode, which can in turn affect the recipient cortex^42,70–72^. One possibility is that neuromodulators utilize strong thalamic connections to influence PNs that track behavioral states, rather than directly driving these neurons (Extended Data Fig. 18)^42,70–72^.

The finding that movement-encoding neurons receive a smaller fraction of inputs from motor cortical areas is somewhat perplexing given the general association of motor cortical areas with movement execution. Motor cortical areas have been shown to be required for learning and production of skilled and accurate movements^73^, rather than for the execution of innate movement sequences. Instead of being learned motor skills, spontaneous whisking and locomotion accompanied by rhythmic whisking are innate behaviors and part of the whole range of coordinated movements that represents a behavioral state. While inactivation of wM1 affects wS1 activity, it does not abolish the behavioral state-dependent changes in S1 activity^74,75^. M1/2 axons in wS1 do not appear to be exclusively dedicated to transmitting movement features, but instead carry multiple aspects of sensorimotor behavior including touch^28^. The M1/2 pathway is known to alter cortical sensory responses through the engagement of a local disinhibitory circuit that changes stimulus response gain to relevant inputs^49,76^. One possibility is that the M1/2 pathway may be more relevant for sensorimotor learning in wS1, instead of determining behavioral state-dependent representations. If the role of these behavioral state-encoding neurons is to reliably update the behavioral state to the local network, one might expect these neurons to be less susceptible to the plasticity of sensorimotor learning.

We employed a single-cell-based monosynaptic retrograde tracing approach using a modified rabies virus, which is a powerful method for mapping brain-wide presynaptic inputs. However, this approach is limited due to only a fraction of inputs being labeled, lack of knowledge regarding the strength of labeled connections, and potential bias associated with viral tropism. To overcome these issues, we directly compared the brain-wide presynaptic networks of two functionally distinct groups using the same method. If a bias exists, it is reasonable to assume that it is similar in both groups. Our study is based on the proportion of presynaptic cells, in different brain areas, of individual cortical neurons. While we focused on the functional relevance of the most salient input pathway revealed by our presynaptic network study (thalamic), other input pathways may also play a significant role in driving postsynaptic activity. It is undetermined how the presynaptic proportions are related to the synaptic strength of inputs from a given brain area, whether the presynaptic cells are functionally similar, and how connectivity relates to the intrinsic properties of postsynaptic neurons.

We found highly convergent yet characteristic presynaptic connectivity patterns depending on the functional property of each neuron. This raises questions about the functions and projection patterns of the movement-correlated neurons within the local network, and their connections to other brain areas. These neurons may report moment-to-moment behavioral states within the sensory cortex and provide a self-referenced framework for integrating information across other brain areas. The stable, sensory-independent activity of wS1 L2/3 neurons may constitute a reflection of internal models of the body and environment that allow for the integration of sensory information with external and internal contexts, prediction of sensory consequences, and guidance of actions^77^. Parallel to their input connectivity, their output connectivity may follow similar rules, whereby functionally distinct neurons project to a wide range of other cortical areas, yet with a selective amplification or reduction of specific projections^78,79^. This information will be critical to build a full model of the anatomical and functional landscape for heterogenous cortical neurons.

The enhanced thalamic input fraction of neurons that tracked behavioral state may arise during development, as suggested by the fact that early cortical activity is dominated by thalamic inputs triggered by spontaneous movements^80–83^. Whether the enhanced thalamic fraction of movement-encoding neurons observed in adults constitutes a developmental trace remains to be explored. Our results revealed anatomical biases in long-range glutamatergic inputs mapping onto functionally distinct neurons. These specific input patterns together with the stable representation uncovered here suggest the existence of preconfigured patterns of activity in S1 in the context of behavioral state^77^.

## Methods

### Mice

All animal procedures were conducted in accordance with a protocol approved by the National Institutes of Health Animal Care and Use Committee and complied with Public Health Service policy on the humane care and use of laboratory animals. We used the following transgenic mouse lines: Emx1-IRES-Cre (JAX 005628)^84^, tetO-GCaMP6s (JAX 024742)^85^, and CaMK2a-tTA (JAX 007004)^86^. We performed experiments on 10 Emx1-IRES-Cre, 71 tetO-GCaMP6s;CaMK2a-tTA, and 3 GCaMP6s^+^.CaMK2atTA^-^ mice. We used both male and female mice (females, 45%), 12-24 weeks old at the experimental endpoint. The percentage of Mov_up_ and Mov_down_ neurons and sensory-responsive neurons was similar between the male and female groups (Mov_up_ and Mov_down_ neurons, 30 ± 11% vs. 33 ± 11 %, *n* = 26 vs. 21 animals, respectively, *P* = 0.27; sensory-responsive neurons, 13 ± 5.9% vs. 12 ± 6.1%, *n* = 19 vs. 14 animals, respectively; *P* = 0.57, Wilcoxon rank-sum test). Mice were housed in groups, in individually ventilated and enriched laboratory cages, in climate-controlled rooms (T, 22 °C; humidity, 45%), under a reverse 12 h light - 12 h dark cycle (light on, 9 a.m.), and with *ad libitum* access to water and food. After surgical procedures, mice were housed individually. All experiments were performed in the dark phase of the cycle. Animals in test and control groups were littermates and randomly selected.

### Surgeries

All surgical procedures were performed stereotaxically, including injection of recombinant adeno-associated viruses (rAAV), head plate implantation, and cranial window implantation, and were carried out under aseptic conditions. Mice were anesthetized with isoflurane (1.0 - 2.0% in O_2_ at 0.8 L/min). The eyes were protected with ophthalmological ointment, and body temperature was maintained at ∼37 °C using a heating pad (Stoelting). Dexamethasone (0.2 mg/Kg of body weight; subcutaneous injection) was administered at least 1 h prior to cranial window implantation, to prevent brain edema. Exposed dura mater was perfused with sterile Ringer’s solution (in mM, 150 NaCl, 2.5 KCl, 10 HEPES, 2 CaCl_2_, 1 MgCl_2_; pH 7.3 adjusted with NaOH; 300 mOsm). After surgery, mice were treated with meloxicam (2 mg/Kg; subcutaneous injection) every 24 h for 3 days, to minimize pain and inflammation, and with enrofloxacin (0.1 mg/mL in drinking water) for 5 to 10 days, to prevent infection. Wellness and body weight were monitored daily for 10 days.

The first surgery consisted of rAAV injection combined with headpost and cranial window implantations, rAAV injection followed by headpost implantation, or headpost implantation only. For rAAV injection, at each target coordinate, the skull was thinned, and a craniotomy (∼50 µm diameter) was made using fine forceps. A glass micropipette (5-10 µm outer diameter tip) attached to a nanoinjector (WPI) was used to deliver the viral vector (at 20-50 nL/min). After injection, the pipette was left in place for ∼5 min before being slowly retracted. To express GCaMP6f in PNs, we injected rAAV5-Syn-Flex-GCaMP6f-WPRE-SV40 (Addgene 100833) in the right hemisphere wS1 of Emx1-IRES-Cre mice (in mm relative to Bregma: AP -0.80 and ML 3.50; AP -1.20 and ML 3.40, the pipette was angled at 21 °, and 30 nL were injected at the subdural depths of 350, 250 and 150 µm). For GRAB sensor experiments, we injected AAV9-hsyn-Ach4.3 (Ach3.0) or AAV9-hsyn-NE2m (NE3.1) (WZ Biosciences) in the right hemisphere wS1 of GCaMP6s^+^.CaMK2atTA^-^mice (3 injection sites; same injection parameters as above). For simultaneous imaging of L2/3 PNs and optogenetic suppression of thalamic or motor cortical terminals in wS1, we injected rAAV5-CAG-ArchT-tdTomato (UNC Vector Core AV4595B), rAAV5-CAG-tdTomato (Addgene 59462), or rAAV5-Syn-tdTomato (Addgene 51506) in the VPm (AP -1.70, ML 1.85, and DV 3.15; 50-60 nL) and/or POm (AP 2.00, ML 1.32, and DV 3.00; 50-60 nL), or the M1/2 (AP -1.00, ML 1.00, and DV 0.8 to 0.2, 15 nL per each 100 µm) of tetO-GCaMP6s;CaMK2a-tTA mice. We implanted a custom-made Y-shaped titanium head plate using dental cement (Super-Bond C&B, Parkell). The exposed skull was covered with a thin layer of clear dental cement and, subsequently, opaque biocompatible silicone (Kwik-Cast, WPI), if applicable. A craniotomy was made over the right hemisphere wS1 (centered at AP 1.1 and ML 3.3), and a glass cranial window (diameter, 3 mm; thickness, 100-150 µm) was placed and secured over the craniotomy using cyanoacrylate adhesive (3M) and dental cement. For in vivo neuropharmacological experiments and single neuron monosynaptic input tracing, the cranial window had a rectangular laser-cut opening (0.30 x 0.80 mm, Potomac Photonics) covered with transparent biocompatible silicone (Kwik-Sil, WPI)^87^. On the day of imaging, the silicone plug was removed and micro-durotomy was performed for direct access to the brain.

Recordings were initiated after a minimum period of 3-5 weeks post-rAAV injection, for stable expression of GCaMP6f, ArchT, GRAB_ACh_ or GRAB_NE_.

### Behavior

All behavioral experiments were performed in darkness. Under head-fixation, mice with all intact whiskers were free to run on a wheel. The mouse face and whiskers were video-recorded at 250 fps using a high-speed camera (acA2000-340kmNIR, Basler) with an 8 mm lens (LM8JC, Kowa), under infrared LED illumination (850 nm, Mightex). Image acquisition was controlled by StreamPix (NorPix). The wheel (diameter, 15 cm; width; 5.5 cm) was set so that only forward movement was permitted. To extract locomotion speed, we used a 2500 CPR resolution motion encoder (Model 260 Accu-Coder, Encoder) affixed to the wheel shaft. Motion encoder pulses were converted to speed using a counter and LabView software (National Instruments (NI)), for online visualization of speed. To synchronize behavioral and neuronal data, voltage signals from each video-frame exposure and wheel speed were digitized and recorded at 10 kHz through a data acquisition card (PCI-6052E, NI), using the Prairie View Interface (Bruker).

### Unilateral mystacial pad paralysis

In a subset of animals, paralysis of the left or both mystacial pad(s) was achieved through a local, subcutaneous injection of BTX (single injection per pad; 0.5 Units in 10µL per injection, BOTOX), under isoflurane anesthesia.

### Whisker stimulation

Deflection of the left whiskers was achieved using a solenoid valve-controlled pole (diameter, 3 mm; length, 5 cm). To maximize contact with all whiskers, the pole was positioned in alignment with the mystacial pad, at an angle of ∼65 ° (distance between the mystacial pad and the pole during stimulation, ∼5 mm). Stimuli (trains of 28-49 stimuli; speed, ∼600 mm/s; duration, 50 or 250 ms; interval, 3-5 s) were produced and synchronized to behavioral and neuronal data using the Prairie View Interface (Bruker). To ensure that changes in neuronal activity related to whisker deflection could be isolated from changes in neuronal activity related to potential alterations in movements during deflection, recordings included trials of whisker deflections coupled with sound, as well as sound-only trials^61^. Neurons were classified as sensory stimulus-responsive if they responded exclusively during whisker deflections, but not during sound-only trials. The percentage of sensory-responsive neurons did not differ when using 50 ms-duration vs. 250 ms-duration whisker stimuli (14 ± 5.9% vs. 11 ± 6.0%, *n* = 21 vs. 12 animals, respectively; *P* = 0.15, Wilcoxon rank-sum test).

### In vivo imaging and optogenetics

Imaging was performed using a two-photon microscope (Ultima Investigator, Bruker) and a fs-pulse Ti:Sapphire laser (Mai Tai DeepSee, Spectra-Physics), tuned between 860 and 1040nm, for imaging of difference fluorescent proteins. The microscope was equipped with an 8 kHz resonant galvanometer and a water-immersion 16X objective (0.8 NA, Nikon) coupled to a 400 µm-range, Z-axis piezoelectric drive. GCaMP6 and RFP fluorescence signals were passed through a 525/70m or 595/50m filter, respectively. Fluorescence was detected and amplified using GaAsP PMTs (Hamamatsu) and a dual preamplifier, prior to digitization. For 2-PT Ca^2+^ imaging, we performed one session per day (recording time ∼66 ± 21 min). Images (resolution, 512 x 512) were collected at ∼30 Hz, in single plane mode. FOVs ranged from 271 x 271 µm (for functional identification of a neuron for subsequent electroporation) to 573 x 573 µm (for characterization of neuronal patterns of activity) with an average excitation laser power of ∼33-76 mW at the objective. For simultaneous 2-PT Ca^2+^ imaging and optogenetic inhibition of thalamic axon terminals, we used a collimated 625 nm LED beam (Prizmatix UHP-T-625-SR)^88,89^. The LED was on during the turnaround of the resonant galvanometer. Trains of light pulses (25 pulses; duration per pulse,1-1.5 s; interval, 5 s) were generated and synchronized to behavioral and neuronal data acquisition using the Prairie View Interface (Bruker). A filter (FF02-617/73-25, Semrock) was used to narrow the LED spectrum, and a dichroic mirror (FF556-SDi01, Semrock) was used direct the LED light onto the brain tissue and pass GCaMP6 florescence signals onto the PMT. The average power of the LED was 10-50 mW at the objective.

### In vivo neuropharmacology

A durotomy (∼50 µm) was made through the access port of the implanted cranial window, under isoflurane anesthesia (Fig. 2a). Mice were allowed to recover for ∼30 min prior to recordings. Following acquisition of baseline behavioral and neuronal activity data, Ringer’s solution was replaced by Ringer’s solution supplemented with receptor blockers, and recordings were reinstated. In sham sessions, Ringer’s solution was replaced by Ringer’s solution without blocker addition. Only data acquired 20 min or more after blocker application or Ringer’s replacement (in sham sessions) were considered for analysis. Each session, 1-2 days apart, lasted a median of 2 h 15 min (effective spontaneous activity recording time,1 h and 8 min): baseline period, 42 min (effective spontaneous activity recording time, 30 min); blocker application or Ringer’s replacement, 3 min; waiting period following blocker application or Ringer’s replacement, 21 min; receptor blockade period, 45 min (effective spontaneous activity recording time, 30 min). Baseline vs. sham/receptor blockade recording times did not differ (*P* > 0.05, Wilcoxon signed-rank test). We used a combination of atropine (1 mM) and mecamylamine (1 mM) to block ACh R^48,49,90,91^, a combination of prazosin (1 mM) and propranolol (1mM) to block NE R^30^, D-AP5 (1 mM) to block NMDA R, and a combination of D-AP5 and DNQX (2 mM) to block both NMDA and AMPA R. In one mouse, we used prazosin (1 mM), propranolol (1mM) and yohimbine (1 mM) to block NE R. Blocker application session sequences were either ACh R – NE R – NMDA/Glut R (*n* = 3) or NE R – NMDA/Glut R – ACh R (*n* = 2), randomly assigned per animal. In two mice, only ACh R (*n* = 1) or NE R (*n* = 1) blocker sessions were performed. The position of ACh or NE R in the sequence did not affect neuronal correlations (*P* > 0.05, linear mixed model controlling for days as confounding factor for position). To test the effectiveness of drug diffusion into the imaging FOV (481.4 x 481.4 to 572.9 x 572.9 µm^2^, at 330 ± 30 µm of depth), we applied TTX (10-100 µM) in the final recording session. Experiments in which TTX did not silence neuronal activity over the entire FOV within 15-20 min were excluded (*n* = 2 mice).

To evaluate the effectiveness of ACh and NE R blockade throughout the entire FOV, we performed equivalent neuropharmacological experiments in mice expressing either GRAB_ACh_ or GRAB_NE_ in wS1 L2/3 neurons.

### In vivo single neuron monosynaptic input tracing

After micro-durotomy, we performed 2-PT Ca^2+^ imaging and selected a target neuron based on its activity profile across behavioral states. Classification of the target neuron as movement-uncorrelated or movement-correlated was confirmed during post-hoc analysis. After imaging, the mouse was lightly anesthetized, and 2-PT guided electroporation of the target neuron was performed as described previously^16,19,50^. A glass pipette (14 ± 1.5 MΩ) was filled with intracellular solution (in mM, 130 K-gluconate, 6.3 KCl, 0.5 EGTA, 10 HEPES, 5 Na_2_-phosphocreatine, 4 Mg-ATP, 0.3 Na-GTP; pH 7.4 adjusted with KOH; 280-300 mOsm) supplemented with Alexa 594 hydrazide (50 µM, A10442, Thermo Fisher Scientific) and two DNA plasmids (pAAV-EF1α-mTagBFP-HA-T2A-mCherry-TVA-E2A-N2c, 0.15 µg/µL; pAAV-CAG-N2c, 0.05 µg/µL). The resistance of the pipette tip was monitored continuously (Axoporator 800A, Molecular Devices). Positive pressure was applied to the pipette (70 mBar), which was visually advanced through the durotomy, using a micromanipulator (PatchStar, Scientifica). Upon entering the cortex, the pressure was swiftly decreased (35 mBar). Then, within ∼50-100 µm from the target neuron the pressure was further decreased (15 mBar). The pipette was slowly advanced towards the soma of the target neuron, until the tip resistance increased by at least 20%. The pressure was released, and a train (100 Hz, 1 s) of electric pulses (-10 V, 0.5 ms) was applied (Axoporator 800A), after which the pipette was retracted. The electroporated neuron was imaged 20 min later to evaluate its survival. Thereafter, we injected G-deleted, envelope-A coated CVS-N2c RV carrying RFP (kindly provided by the Center for Neuroanatomy with Neurotropic Viruses) in the vicinity (within ∼150 µm) the electroporated neuron (rate, 30 nL/min)^92^. Then, the access port of the cranial window was sealed using biocompatible silicone. Survival and successful transfection of the electroporated neuron was monitored within 2-3 days after electroporation and up to the experimental endpoint. Structural, 2-PT z-stacks (1-5 µm-steps; resolution, 512 x 512; FOV, 102 x 102 to 271 x 271 µm) including the imaging FOV and/or the target neuron were acquired before and after electroporation, to track individual cells volumetrically throughout the experiment. Local, wS1 presynaptic networks were followed structurally through 2-PT imaging of GCaMP6 and RFP (z-stacks, 1-5 µm-steps; resolution, 512 x 512; FOV, 271 x 271 to 814 x 814 µm). For z-stack acquisition the average laser power was depth-adjusted linearly, at did not exceed 100-150 mW at the objective. Mice were sacrificed at day 11 (+/- 1.5 days) following electroporation, and brains were processed for ex vivo input tracing. Brains containing less than 100 presynaptic cells were excluded from analysis (*n* = 1).

### Histology

Upon completion of recordings, mice were deeply anesthetized and perfused transcardially with 4% formaldehyde in PBS. Post-perfusion, brains were immersion-fixed in 4% formaldehyde in PBS for 2-3 h and then transferred to 30 % sucrose in PBS.

For input tracing experiments, whole brain free-floating sequential coronal sections (50 µm-thick) were obtained using a microtome (SM2010R, Leica). Sections were rinsed 3 times in PBS, incubated in blocking solution (5% normal serum and 1% TritonX-100 in PBS) at room temperature (RT) for 1 h, and subsequently incubated in primary antibody solution (2% normal serum and 1% TritonX-100 in PBS) at 4 °C for 48 h. Primary antibodies were detected through incubation in secondary antibody solution (2% normal serum and 1% TritonX-100 in PBS) at RT for 2 h. We used the following primary and secondary antibodies and respective dilutions: anti-RFP 1:500 (600-901-379, Rockland); anti-GABA 1:500 (A2052, Sigma), IgY-Alexa Fluor 555 1:200 (A21437, Thermo Fisher Scientific); IgG-Alexa Fluor 647 1:200 (A21245, Thermo Fisher Scientific). Sections were rinsed in PBS and sequentially mounted on glass slides. Neuronal nuclei were revealed through a fluorescent Nissl stain (NeuroTrace 435/455, N21479, Thermo Fisher Scientific), after which sections were cover-slipped. Whole-brain serial sections were imaged using an epifluorescence illumination microscope (Axio Imager, Zeiss). Multiple z-stacks (10 µm steps), covering each section in its entirety, were acquired using Neurolucida (MBF Bioscience). Z-stacks were aligned and collapsed onto a single image using Deep Focus (Neurolucida).

For all other experiments, brain sections were similarly generated and mounted, and neuronal nuclei were visualized either using fluorescent Nissl stain (NeuroTrace 435/455 or NeuroTrace 530/615, N21482, Thermo Fisher Scientific) or DAPI (Fluoromount-G mounting medium, Thermo Fisher Scientific). Entire sections were imaged using an Axio1 Scanner (Zeiss). FOV location within wS1 was confirmed either by targeted 2-PT laser microlesions (the laser beam was focused at a subdural depth of 200 µm; 800 nm; 5-30 s; ∼0.5 W)^93^ or fluorescent die (DiI (42364, Sigma) or Fast Blue (17740, Polysciences)) injection at the experimental endpoint and ex vivo histological analysis. Tissue imaging and histological analysis were done blinded to the experimental groups. For analysis of thalamic ArchT-expression areas, a composite image of a brain section at the injection center for either VPm or POm was selected, and expression areas annotated; each brain section was manually aligned to the corresponding mouse brain atlas section, and the expression areas of the different animals were overlayed^94,95^.

### Defining behavioral events

Video recordings of face and whiskers were processed using a custom MATLAB routine. To detect whisker movements, we first defined a region of interest (ROI) encompassing the left or right whiskers in an image that consisted of the s.d. of representative frames of each session. We then computed the absolute power of the spatial derivative of consecutive frames (whisk. mov. trace). Both the whisker movement and locomotion speed traces were downsampled to 30 Hz, averaging the values acquired during a Ca^2+^ imaging frame. To detect behavioral events, the whisker movement trace was baseline subtracted (10^th^ percentile of the full trace) and normalized. We then applied a threshold to the whisker movement trace (3 x minimum s.d., calculated using a 30 s sliding window). Detection was visually inspected for all sessions, and the threshold was adjusted when applicable. Then, local maxima were calculated, and peaks less than ∼0.5 s apart were considered as part of a single event; the first peak was considered as onset and the last, as offset. Events with an integral value smaller than 20 (< 3% of the session time) were excluded from analysis. Whisker movement events were considered as WL when the maximum locomotion speed was higher than 0.20 cm s^-1^ and, conversely, as W_only_ when the locomotion speed did not exceed 0.20 cm s^-1^. For comparisons across animals, the raw whisker movement trace was baseline subtracted and normalized to its maximum; for BTX experiments, across session data were normalized to maximum according to the first session.

### Processing of two-photon calcium images

Two-photon Ca^2+^ images were processed using Suite2p^96^, in Python, with default parameters, unless otherwise indicated. Following subtraction of neuropil (fixed scaling factor of 0.7) and baseline (calculated on filtered traces, using a gaussian kernel of width 20 and a sliding window of 60 s), fluorescence traces were deconvolved using non-negative spike deconvolution^97^ with a fixed decay timescale of 0.7 s for GCaMP6f and 1.5 s for GCaMP6s. To make sure that only somatic traces were included in the analysis, ROIs were manually curated by an analysist blinded to the experimental group. Aligned image series were visually inspected to control for z-drifts; data showing z-drifts were excluded from analysis. Tracking of the same neurons across sessions was done semiautomatically using a MATLAB script. All analysis was based on deconvolved traces; for presentation purposes only, we used neuropil subtracted fluorescence traces normalized to the maximum (F), overlayed with deconvolved fluorescence traces normalized to the maximum (F deconv.), unless otherwise indicated. To generate temporal raster plots (Fig. 1e), the activity of each neuron was averaged over ∼0.5 bins, z-scored and smoothed using a 1 s moving average filter; individual neurons were sorted by the first principal component of neuronal activity.

### Processing of two-photon GRAB sensor images

Two-photon Ca^2+^ images were motion corrected using Suite2p^81^. Thereafter, we extracted the mean fluorescence intensity of pixels within six ROIs (75 x 75 pixels) manually spread over the entire FOV, avoiding large vessels. In addition, to extract a full FOV fluorescence intensity mean, we first isolated sensor^+^ pixels and excluded vessel-related pixels by applying a threshold to pixel intensity (99.9^th^ percentile < intensity > 50^th^ percentile) over the motion-corrected, whole recording session average image^98^. Baseline traces were obtained essentially as described above (Processing of two-photon Ca^2+^ images) but using a gaussian kernel of width 30 and a sliding window of ∼120 s. The mean fluorescent values of single ROIs or full FOVs were baseline subtracted (ΔF) and z-scored.

### Decoding analysis

We trained a linear decoder to decode behavioral variables from neuronal activity. We minimized the ridge regression^99^ objective function (Eq. 1)

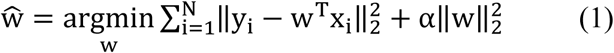

where y_i_ is the behavioral variable at time (frame) i, x_i_ is the neuronal activity matrix, w is the weight vector, and α is the ridge parameter (regularization). We normalized the behavior and neuronal activity by z-score for the cross-day recordings. We did not normalize the neural activity for the neuromodulatory experiments, as activity was recorded continuously on the same day. For optogenetic experiments, we ran analysis with (shown in figures) and without data normalization, as well as with and without (shown in the figures) rebound cells, and no substantial difference was found. We randomly split the trials into training (75%) and test sets (25%). The weight vector was estimated on the training set, and the ridge parameter was selected by leave-one-out (LOO) cross-validation^100^ on the training set.

We evaluated the decoding performance by the out-of-sample (test set) coefficient of determination (R^2^) (Eq. 2)^99^

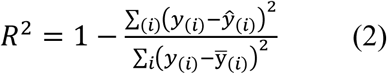

where (i) is the index of out-of-sample trials, ŷ_(i)_ = ŵ^T^x_(i)_ and ŵ is the estimated weight vector from the training set by (1). Using the weight vector estimated from the training set, we decoded the behavioral variables on the test (held out) set within the same condition or session (referred as within) and the trials of other conditions or sessions (referred as across). Then, we calculated the out-of-sample R^2^ using the predicted and true values for within and across data. The out-of-sample R^2^ is not prone to overfitting and will not be inflated. Note that the out-of-sample (test set) R^2^ can be negative if the fitted model does not predict the test set at all, which indicates that the neural correlations could be substantially different between the training and test sets.

### Modulation of neuronal activity

Data were analyzed using MATLAB scripts. The activity (F deconvolved) of each neuron was aligned to the onset and offset of spontaneous movements, W_only_ and WL. Baseline and post-offset activities refer to a 0.5 s window preceding movement onset and a 0.5 s window after movement offset, respectively. A neuron was considered as Mov_up_ if its average activity during W_only_ and/or WL events was significantly higher than its average activity during baseline and/or significantly higher than its average activity post-event (*P* < 0.01, paired-sample *t*-test). Conversely, a neuron was considered Mov_down_ if its average activity during W_only_ and/or WL events was significantly lower than its baseline and/or significantly lower than its post-event average activity (*P* < 0.01, paired-sample *t*-test). Neurons exhibiting opposite changes in activity for W_only_ and WL were rare (0.7 ± 0.2% of all cells) and were not considered for further analysis. Otherwise, neurons were considered as movement-uncorrelated. Modulation refers to the mean activity during movement, irrespective of movement duration, minus the mean activity during baseline, averaged across spontaneous movements (W_only_, WL, or W_only_ + WL).

Time-normalized PETHs were created by normalizing the data to match the average duration of WL events across animals. To compute the correlation (Pearson’s linear correlation coefficient, *r*) between the activity of individual neurons and each behavioral variable, deconvolved fluorescent traces, and corresponding whisker movements and locomotion speed raw traces were binned (bin size, ∼0.5 s).

Sensory stimulus-responsive neurons were defined by a significantly higher average activity within a response window of 0.5 s after the onset of whisker stimulation (coupled with sound) vs. baseline (*P* < 0.01 paired-sample *t*-test). We excluded cells that also responded to sound-only stimuli vs. baseline (paired-sample *t*-test, *P* < 0.01). Baseline was calculated on a 0.5 s window preceding stimulus onset. Sensory stimulus-response magnitude refers to the mean activity during the response window minus the mean activity during baseline, averaged across all whisker stimulations.

For neuropharmacological experiments, the distribution of modulation values for each WL neuron in the presence of receptor blocker(s) was compared to that of baseline (prior to blocker application, *t*-test). Similarly, we compared the distribution of sensory stimulus-response magnitude values before and after blocker application for each sensory stimulus-responsive neuron (*t*-test).

For optogenetic experiments, light off periods were used to identify neurons as movement-uncorrelated or movement-correlated. Neurons were classified as light modulated if their activity during the light pulse differed significantly from that of baseline (0.5 s window prior to light pulse onset). Most L2/3 PNs fire sparsely; only neurons that showed an average baseline activity (F. deconvolv.), i.e., prior to the light pulse, higher than the 75^th^ percentile of the average baseline activity across all cells were included in the analysis. To generate time-normalized PETHs, light pulse data were normalized to match the maximum pulse duration across experiments (1.5 s). Statistical testing on PETHs of movement-uncorrelated and movement-correlated neuronal subpopulations was performed on the original data sampling rate (∼30Hz).

### Whole-brain reconstruction, annotation, and registration

To analyze the brain-wide distribution of presynaptic neurons, we adapted a previous pipeline^101,102^. To reconstruct a whole brain 3-dimentionally, individual section images were aligned using BrainMaker (MBF Bioscience). Brain-wide presynaptic neurons (RFP^+^) were automatically segmented using NeuroInfo (MBF Bioscience) and manually annotated according to brain area. Local, wS1 presynaptic neurons were manually annotated based on cortical layer location. Distinct layers were identified based on the characteristic depth-varying density of NeuroTrace^+^ neurons. Presynaptic neurons within each layer were identified as glutamatergic (GABA^-^) or GABAergic (GABA^+^). To confirm the colocalization of RFP and GABA in, a subset of brains was re-imaged using a confocal microscope (z-stacks, 3 µm steps, C2, Nikon). Identification of glutamatergic and GABAergic neurons was equivalent for the two different imaging methods. Each serially reconstructed brain was registered to the Allen Mouse Common Coordinate Framework, and brain-wide presynaptic neurons (RFP^+^) were automatically re-identified according to distinct anatomical structures. Reconstructions and registrations were conducted blindly. Manual identification was independently performed by two analysts (one analysist was blinded to the activity profiles of the postsynaptic neurons). All manual and automatic identifications were coherent.

### Analysis of brain-wide presynaptic networks

#### Fine-scale spatial registration of wS1 presynaptic networks

Postsynaptic neurons did not survive until the experimental endpoint^16^. We used the center of mass of glutamatergic presynaptic networks in L2/3 to estimate the position of the postsynaptic neurons for all 22 subjects^16^, which were then averaged to construct a reference postsynaptic site on the Allen Mouse Common Coordinate Framework. From the brain atlas, we manually marked the boundaries of the cortical surface and performed surface triangulation. Using this triangulated cortical surface, we then estimated the surface’s normal vector that goes through the reference postsynaptic site and serves as the reference normal vector. Individual wS1 presynaptic network of each subject was then rigidly aligned based on the position of the reference postsynaptic site and orientation of the reference normal vector.

#### Layer-by-layer horizontal flat projections

After the fine-scale registration, we concatenated all neurons from both movement-uncorrelated and movement-correlated groups. Then we performed PCA on each layer of the presynaptic networks to obtain the population best-fitted plane. Corresponding neurons from the layer were then projected onto the best-fitted plane. We then performed a rigid parameterization and mapped the neurons to a two-dimensional coordinate system. The resulting parameterization of each layer from every subject with gaussian kernel-density estimation is visualized in Extended Figure 11.

#### Statistical analysis of group-wise spatial distribution differences

To explore whether there was any difference in the spatial pattern of presynaptic networks between movement-uncorrelated and movement-correlated groups, we tested the null hypothesis of no spatial distribution difference between multiple local and long-range anatomically annotated presynaptic neurons of the two groups. The final statistical analysis incorporated Bonferroni correction. We chose the 2-Wasserstein distance function as the test statistics, and we performed a randomization test (*n* = 10,000) to approximate the permutation distribution^103,104^. Due to the variability of total number of presynaptic neurons across brains, we introduced non-uniform sample weights when we computed the 2-Wasserstein distance to avoid any dominated effect from subjects having a large number of presynaptic neurons. Specifically, we first re-weighted every neuron by the inverse of the number of (e.g., layer-wise/whole-wise) presynaptic neurons to ensure an equivalent contribution from subjects with non-empty neuron sets. Second, sample weights from subjects in the same group were concatenated and normalized into a probabilistic mass. This alleviates the imbalance of group-wise total mass if some of the subjects have no neurons detected in specific cortical layers (for instance, GABAergic presynaptic neurons in Layer 6). To aid in the visualization of the spatial spread of presynaptic networks across cortical areas, we generated cortical flat maps. The 3D CCF coordinate points of registered neurons were projected along streamline paths orthogonal to the cortical surface to create a flat map 2D representation that preserves relative spatial position of cortical areas for analysis using tools provided by the Allen Institute^105^.

#### Statistical analysis of group-wise proportions of M1 and M2 presynaptic cells

Motor cortical neurons that project to wS1 are distributed closely along the anatomical border between M1 and M2^106^. To evaluate whether the movement-uncorrelated and movement-correlated groups exhibited an M1 or M2 bias in presynaptic cell proportions, in addition to the fraction of M1 and M2 cells, we computed the shortest 3D Euclidean distance of each M1/2 cell from the anatomical border between M1 and M2 (represented by an open surface mesh in the Allen Mouse Common Coordinate Framework). Distances were assigned with negative or positive values based on whether cells were in M1 or M2, respectively. Given the variability in cell counts across subjects, we implemented a reweighting procedure on the distance distribution, aiming at an equitable contribution of each subject to the group. This involved resampling cells based on a probability assigned to each cell, inversely proportional to the cell count within each subject, resulting in groupwise distribution of distances based on 10,000 resampled cells. Due to a minimal number of M1/2 cells, four animals in the movement-uncorrelated and four animals in the movement-correlated groups were excluded. The observed mean difference was compared to a distribution of mean differences obtained by randomly permuting the group labels and recalculating the weighted mean difference for each permutation. This process was repeated 5,000 times to obtain the permutation distribution.

### Statistics

We did not use statistical methods to predetermine sample size. Data were presented as mean ± s.d. throughout the text. In box plots, the central line represents the median, the box represents the 25^th^ and 75^th^ percentiles, and the whiskers extend to the most extreme data points excluding outliers (larger than 1.5 x the interquartile range); when overlayed with individual datapoints, all datapoints, including outliers, were graphed. Statistical tests used were indicated in the figure legends. All comparisons using two-sample or paired-sample *t*-tests, Wilcoxon rank-sum or signed rank tests were two-sided unless otherwise indicated. Linear fixed effects models (with interaction between drug type and neuronal correlations) were used to compare slopes (Fig. 2). Linear mixed effects models were used to test effect of receptor blocker application order. Bonferroni correction was applied to multiple comparisons unless otherwise indicated. Significance levels are indicated as: NS, *P* ≥ 0.05; *, *P* < 0.05; **, *P* < 0.01; ***, *P* < 0.001. Analytical routines and were established using a subset of the data (training set), and these were applied to entire datasets, whenever applicable to avoid overfitting.

## Data availability

The datasets are available from the corresponding authors upon reasonable request.

## Code availability

The custom codes used for analyses have not been deposited in a public repository but are available from the corresponding authors upon reasonable request.

## Acknowledgements

We thank all members of the S. Lee lab, the C. McBain lab, the T. Petros lab, and the W. Lu lab for helpful discussions; M. Han, J. Qi, W. Zhang for technical assistance with genotyping and histology; R. Paletzki for assistance with processing of whole-brain histological images; C. Scott Gerfen for generation of cortical flat maps; the NIMH Section on Instrumentation (G. Dold, D. Ide, J. Kim, and T. Talbot) and the NIH IDEAS lab (L. Argueta, J. Krynitsky, T. Pohida) for providing custom hardware and software for data acquisition; the NIMH Systems Neuroscience Imaging Resource (J. Kuo, T. Usdin, S. Williams) for assistance with histological analyses; the NIMH Rodent Behavioral Core (Y. Chudasama) for providing surgical setups. We thank the Center for Neuroanatomy with Neurotropic Viruses for kindly providing CVS-N2c RV. We thank C.I. Baker, G. Fishell, R. Khazipov, D. Leopold, E. Merriam, B. Rudy, and P.E. Rueda-Orozco for suggestions on the manuscript. This work was supported by the Intramural Research Program of the National Institute of Mental Health, National Institutes of Health (ZIAMH002959 to S.L.; ZIAMH002497-34 to C.R.G.).

## Author contributions

A.I. and S.L. conceived the study and designed the experiments. A.I. performed the experiments. A.I., K.C.L., Y.Z., F.P., C.R.G. and S.L analyzed the data. A.I. and S.L. wrote the manuscript, with inputs from K.C.L., Y.Z., F.P., and C.R.G..

## Competing interests

The authors declare that no competing interests exist.

---

**Extended Data Fig. 1:**
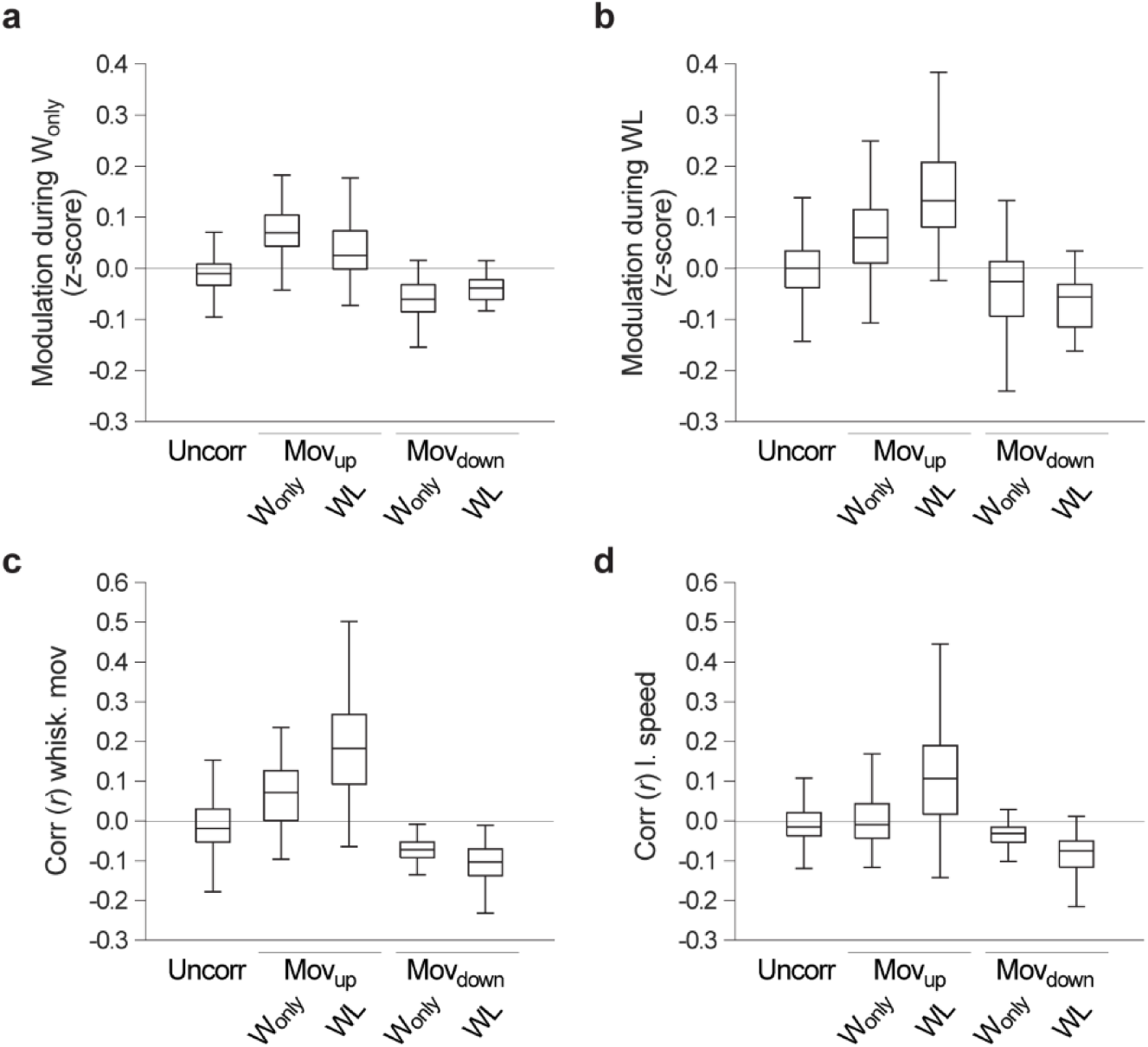
Classification of individual neurons based on their activity during spontaneous movements. **a-d**, To increase neuronal classification accuracy, spontaneous movements were subdivided into two subtypes, W_only_ and WL. Neurons were considered as Mov_up_ or Mov_down_ if their activity changed significantly during either W_only_ or WL, and all other neurons were considered as movement-uncorrelated, resulting in 5 main subsets: movement-uncorrelated, Mov_up_ (W_only_; WL) and Mov_down_ (W_only_; WL). Neurons exhibiting a significant increase or decrease in activity across both spontaneous movement subtypes (W_only_ + WL) were included in the Mov_up_ (WL) or Mov_down_ (WL) subsets, respectively. The proportions of Mov_up_ neurons according to behavioral events were: 25 ± 19% for W_only_, 41 ± 26% for WL, and 34 ± 19% for W_only_ + WL. **a-b**, Modulation (i.e., change in activity during spontaneous movements relative to baseline, prior to movement onset) of neurons classified as movement-uncorrelated (715), Mov_up_ (W_only_, 62; WL, 224), and Mov_down_ (W_only_, 85; WL, 38) during spontaneous movements (6 FOVs, 6 sessions, 5 mice). **a**, Modulation of all subsets during W_only_ (263 ± 98 events per session). Note that Mov_up_ (W_only_) showed the highest modulation. **b**, Modulation of all subsets during WL (50 ± 27 events per session). Note that Mov_up_ (WL) showed the highest modulation (referred to as movement-correlated neurons). Note also that the Mov_up_ (W_only_) cell cluster still shows an increase in activity during WL compared to uncorrelated cells, albeit typically more transient and less robust than that of Mov_up_ (WL). **c-d**, Correlation (*r*) of neurons from the different subsets with whisker movements (**c**) and locomotion speed (**d**). Modulation and correlation values are largely coherent.

**Extended Data Fig. 2:**
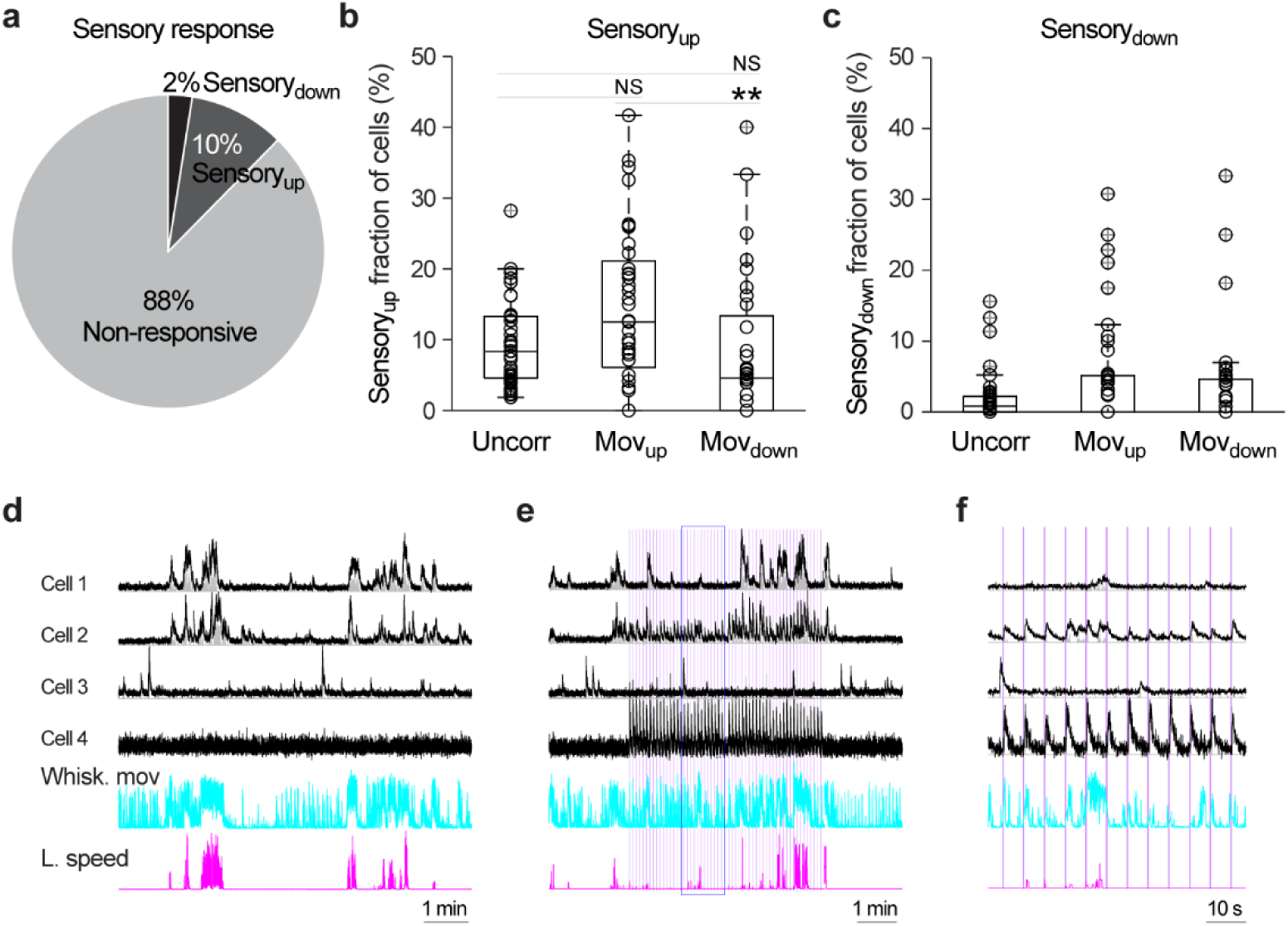
Responsiveness to sensory stimuli. **a**, Fraction of sensory stimulus-responsive neurons from all recorded neurons irrespective of their activity pattern during spontaneous movements. Neurons responded by either increasing their activity (Sensory_up_, 801) or decreasing it (Sensory_down_, 236) during stimulus presentation (*n* = 8750 cells, 44 FOVs, 44 sessions, 33 animals). **b-c**, Fraction of sensory stimulus-responsive neurons within the movement-uncorrelated, Mov_up_, and Mov_down_ subsets. **b**, Sensory_up_ neurons (*P* < 0.01, Kruskal-Wallis test followed by pairwise comparisons). **c**, Sensory_down_ neurons (*P* > 0.05, Kruskal-Wallis test). **d-f**, Example neuronal activity during spontaneous movements and sensory stimulation. Example Mov_up_ (1 and 2) and movement-uncorrelated (3 and 4) neurons that either did not respond (1 and 3) or responded (2 and 4) to sensory stimulation. **d**, Absence of sensory stimulation. **e**, Presence of sensory stimulation. **f**, Expanded time window from **e** (blue box). Black, F; gray, F deconv.; both normalized to maximum. Purple vertical bars, individual sensory stimuli.

**Extended Data Fig. 3:**
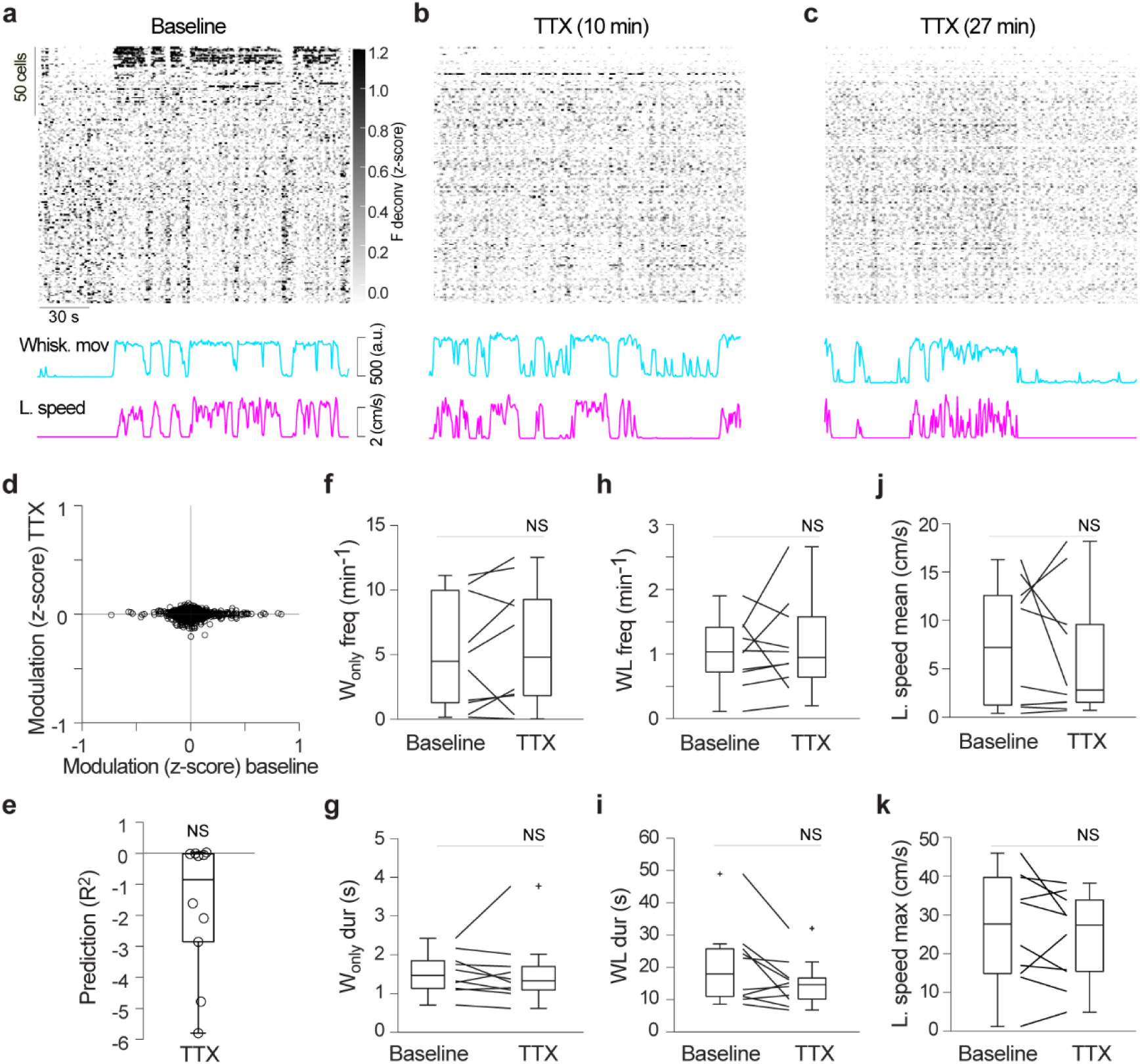
Suppression of cortical activity by TTX diffusion into the brain parenchyma using a custom cranial window. **a-c**, Top, example raster plots of neuronal activity, before and at different time points after application of TTX. Individual neurons (184) are sorted from top to bottom by decreasing weight on the first principal component. Bottom, corresponding whisker movements and locomotion speed traces. **d**, Modulation of individual neurons (1853) during spontaneous movements (W_only_ + WL) before vs. after application of TTX (*P* > 0.05, regression). **e**, Prediction of whisker movements from population activity. Linear decoder predictive R^2^; the decoder was built using baseline data and evaluated on TTX out-of-sample data (*P* > 0.05, paired sample *t*-test across R^2^ ≤ 0 vs. R^2^ > 0). **f-k**, Spontaneous movement parameters before and after application of TTX: W_only_ frequency (**f**), W_only_ duration (**g**), WL frequency (**h**), WL duration (**i**), mean (**j**) and maximum (**k**) locomotion speed (*P* > 0.05 for all panels, Wilcoxon signed-rank test, *n* = 10 sessions, 10 animals).

**Extended Data Fig. 4:**
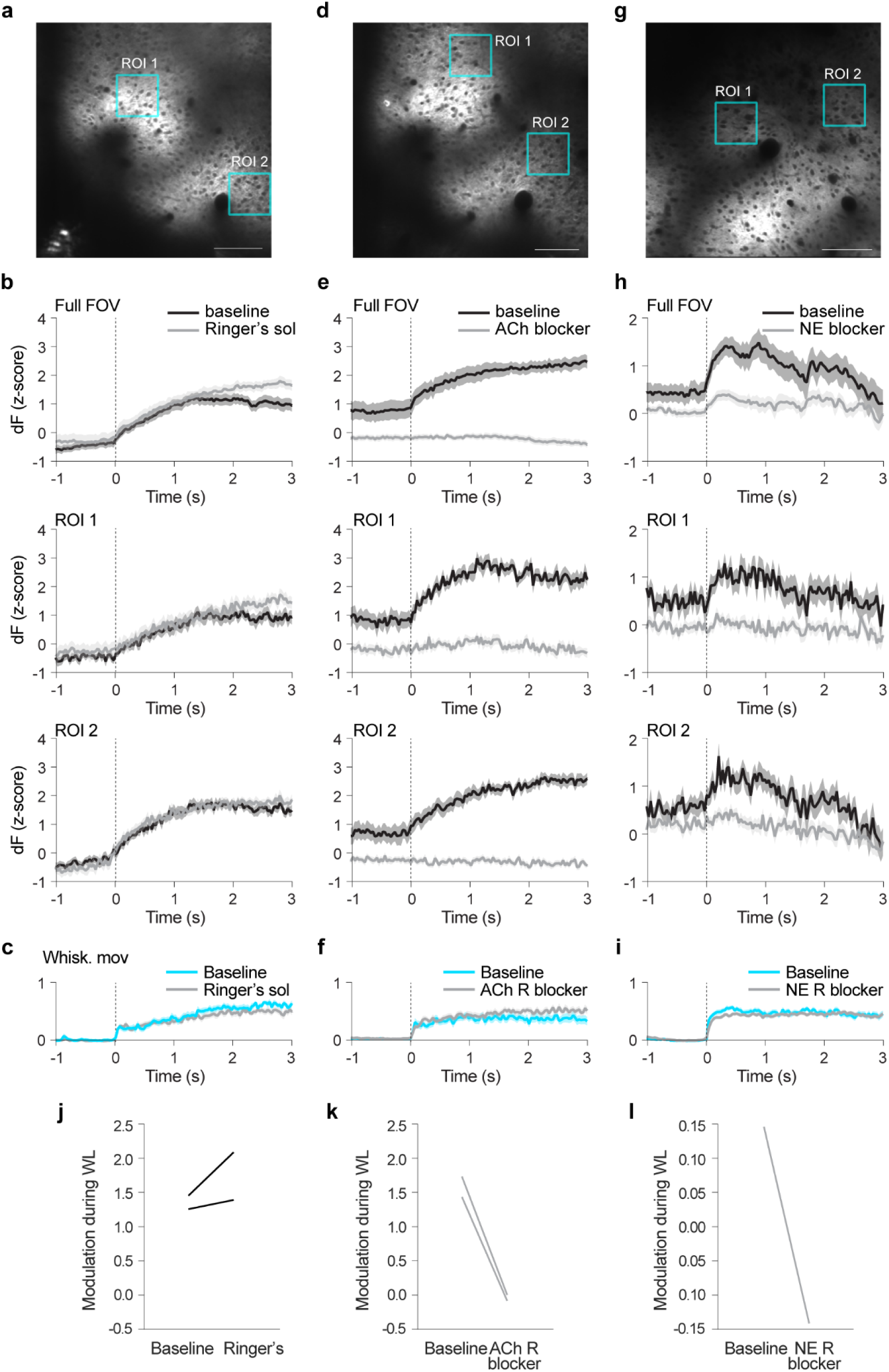
Effectiveness of local application of neuromodulatory receptor blockers through a cranial window port. **a**, Example imaging FOV denoting the expression of the ACh sensor GRAB_ACh_ in wS1 L2/3 neurons. Scale bar, 100 µm. **b**, ΔF averaged over the full FOV and two example ROIs, aligned to the onset of WL events (vertical bars) before and after reapplication of Ringer’s solution in an example mouse (sham session). **c**, Corresponding whisker movements. Results are presented as mean ± s.e.m. **(b-c**). **d-f**, As in a-c, but instead exemplifying the expression of the ACh sensor GRAB_ACh_ and data recorded before and after application of ACh R blockers. **g-i**, As in **a-c**, but instead exemplifying the expression of the NE sensor GRAB_NE_ and data recorded before and after application of NE R blockers. **j-l**, Baseline-subtracted mean ΔF during WL events (modulation during WL) for sham (*n* = 2, **j**), ACh-R blockade (*n* =2, **k**), and NE R blockade (*n* = 1, **l**) sessions. WL-related increases in ACh and NE were abolished when the respective R blockers were applied, but not in sham sessions.

**Extended Data Fig. 5:**
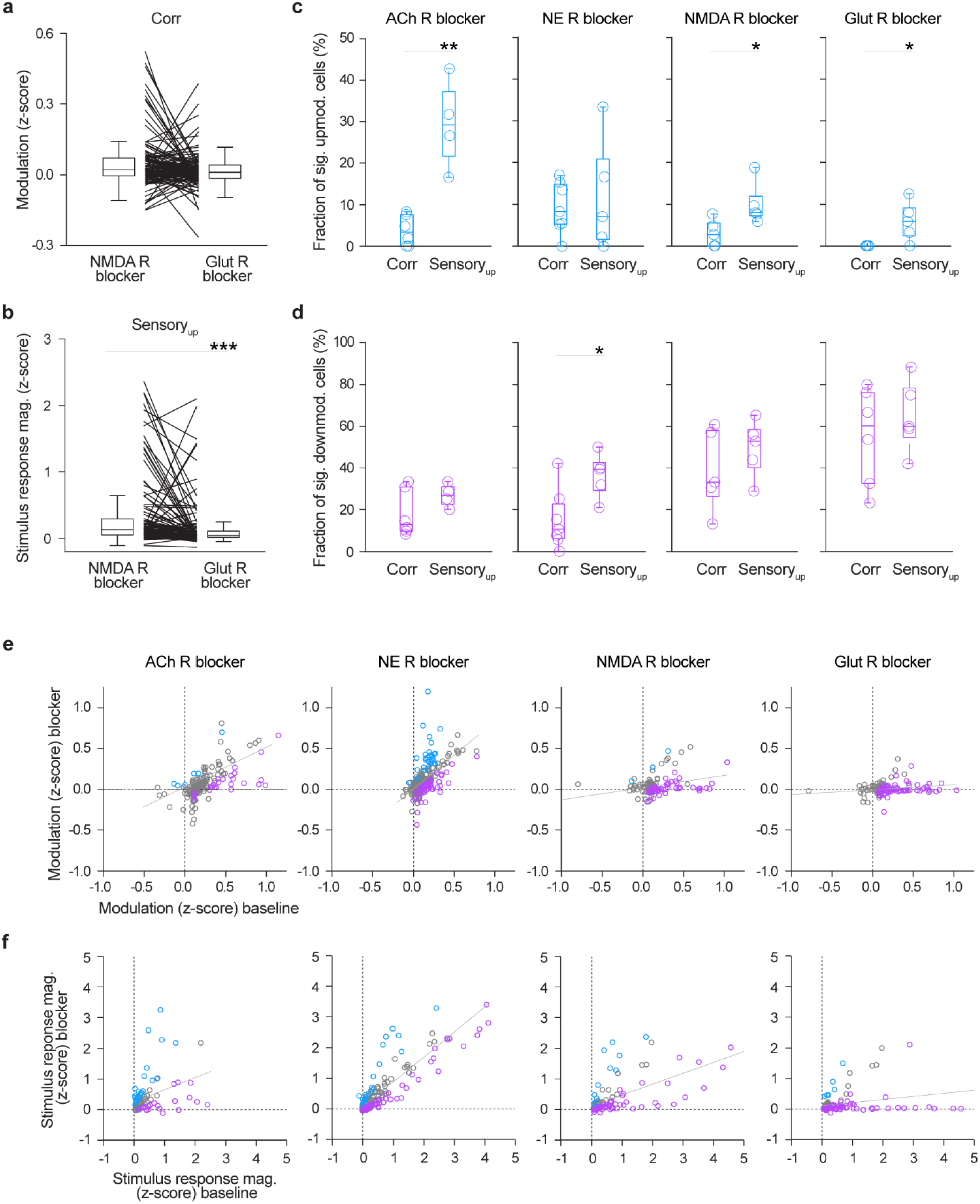
Effect of neuromodulatory inputs on the activity of wS1 L2/3 PNs. **a**, Modulation of individual movement-correlated neurons (118) during spontaneous movements (WL) in the presence of an NMDA R blocker and in the presence of Glut R blockers (*P* > 0.05, Wilcoxon signed-rank test). Neurons were defined as movement-correlated or stimulus-responsive based on data acquired prior to antagonist application. **b**, Sensory stimulus-response magnitude of individual stimulus-responsive neurons (115) in the presence of an NMDA R blocker and in the presence of Glut R blockers (*P* < 0.001, Wilcoxon signed-rank test). Only neurons showing an increased activity following stimulus presentation were included (Sensory_up_). **c**, Fraction of movement-correlated neurons that exhibited a significant increase in modulation during WL, and fraction of Sensory_up_ neurons that exhibited a significant increase in stimulus response magnitude in the presence of either ACh (*n* = 4-6 sessions, 4-6 animals), NE (*n* = 5-7 sessions, 5-7 animals), NMDA (*n* = 5 sessions, 5 animals) or Glut R (*n* = 5-6 sessions, 5-6 animals) blockers vs. baseline (Wilcoxon rank-sum test). **d**, Equivalent to **c**, but for movement-correlated and Sensory_up_ neurons showing a decreased activity. **e**, Modulation of individual movement-correlated neurons (118-273) during spontaneous movements (WL) in the presence of the different blockers vs. baseline (ACh and NE R, *P* < 0.001; NMDA R, *P* < 0.01; Glut R, *P* < 0.05, regression). Purple, cells showing a significant increase. Blue, cells showing significant decrease. **f**, Stimulus response magnitude for individual Sensory_up_ neurons (84-197) in the presence of the different blockers vs. baseline (ACh, NE, and NMDA R, *P* < 0.001; Glut R, *P* < 0.01, regression). The effect of neuromodulatory input blockade, in particular ACh, is more robust is at the level of responses to sensory stimuli.

**Extended Data Fig. 6:**
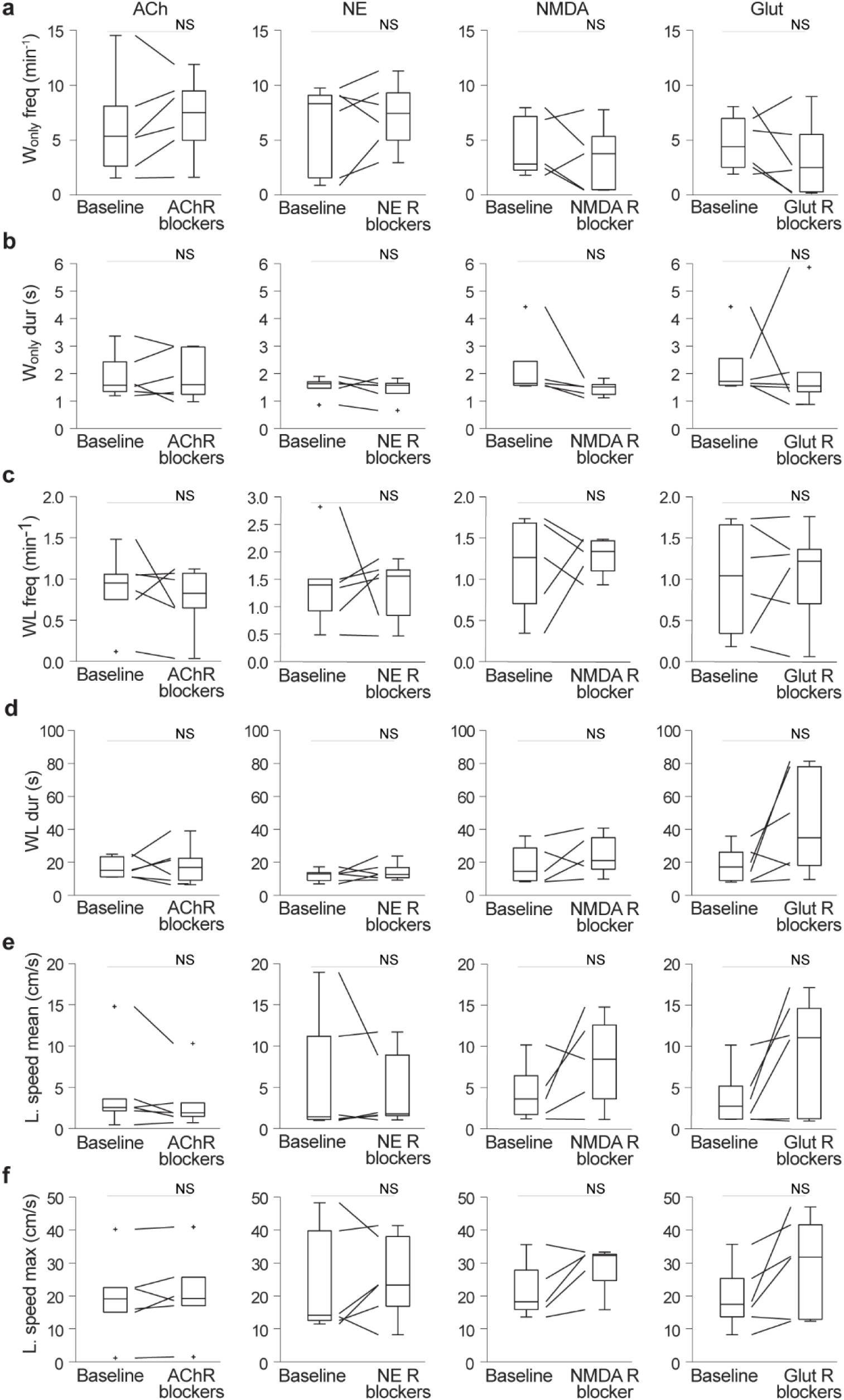
Effect of neuromodulatory and glutamatergic receptor blockade on spontaneous movements. **a-f**, Spontaneous movement parameters, W_only_ frequency (**a**), W_only_ duration (**b**), WL frequency (**c**), WL duration (**d**), mean (**e**) and maximum (**f**) locomotion speed prior to (baseline) and during ACh, NE, NMDA or Glut R blockade (*P* > 0.05 for all panels, Wilcoxon signed-rank test, *n* = 5-6 sessions, 5-6 animals).

**Extended Data Fig. 7:**
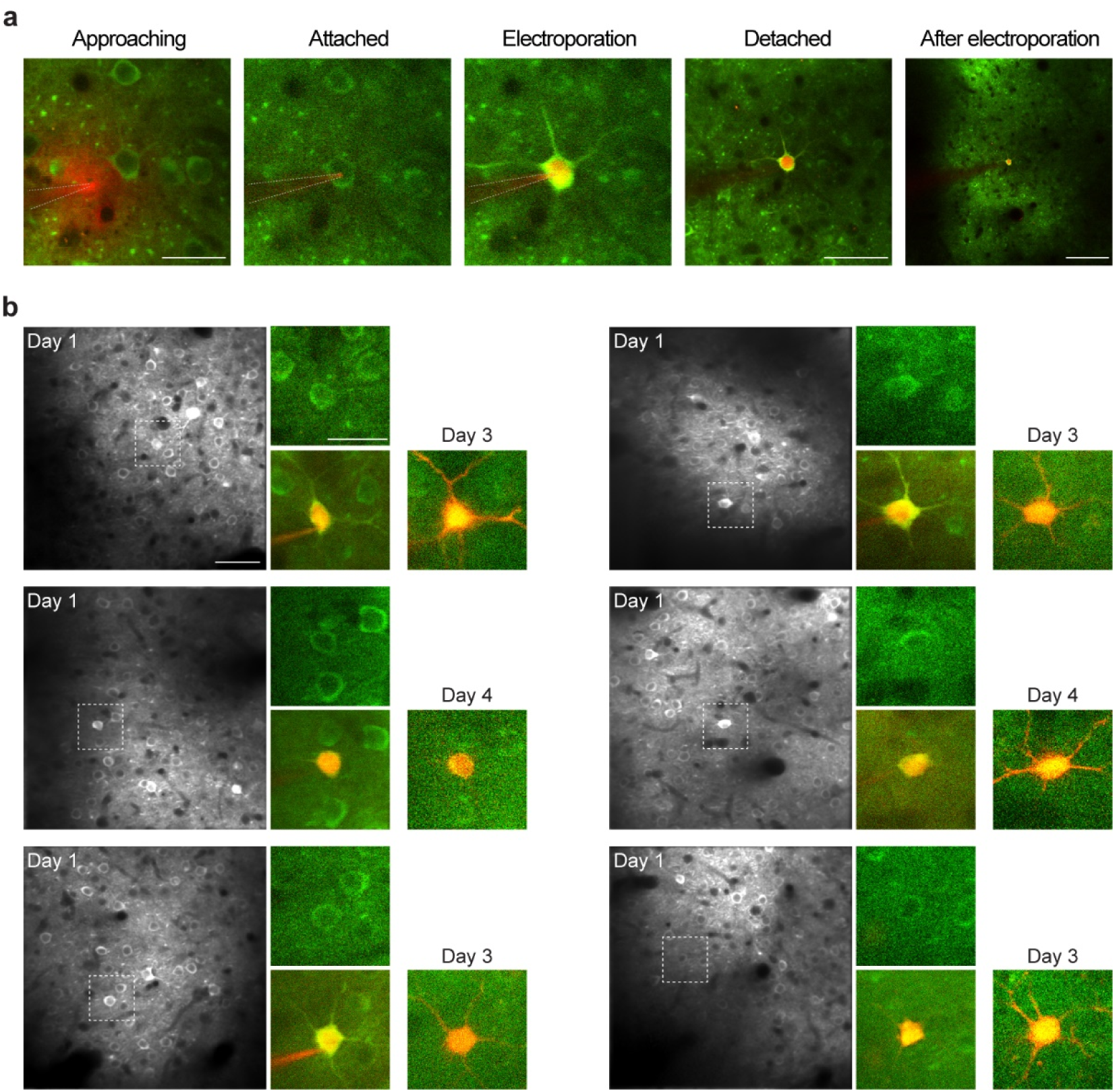
Electroporation of a functionally defined single neuron per brain. **a**, Left to right, Example 2-PT guided approach of a PN in wS1 L2/3 with a pipette containing intracellular solution, Alexa 594 (red) and DNA (for TVA, G, and mCherry). Principal neurons express GCaMP6 (green). The 2-PT laser excitation wavelength is 860 nm, which allows imaging both GCaMP6 and Alexa 594. The pipette tip is advanced against the cell body until resistance increases by at least 20%, and the positive pressure applied to the pipette is then gently released. Once the pipette tip establishes stable contact with the cell body, an electric pulse is applied, for electroporation, followed by gentle retraction of the pipette. Electroporation is indicated by the entry of Alexa 594 into the target cell. Images of the brain and target neuron at different magnifications. Scale bars, from left to right, 25, 50, and 100 µm. **b**, Example postsynaptic neurons. Left, Imaging FOV. Scale bar, 50 µm. Middle, high magnification 2-PT images of the postsynaptic neuron before and immediately after (detached pipette) electroporation. Right, Image of the target neuron 3-4 days after electroporation (GCaMP6s^+^/mCherry^+^). Scale bar, 25 µm.

**Extended Data Fig. 8:**
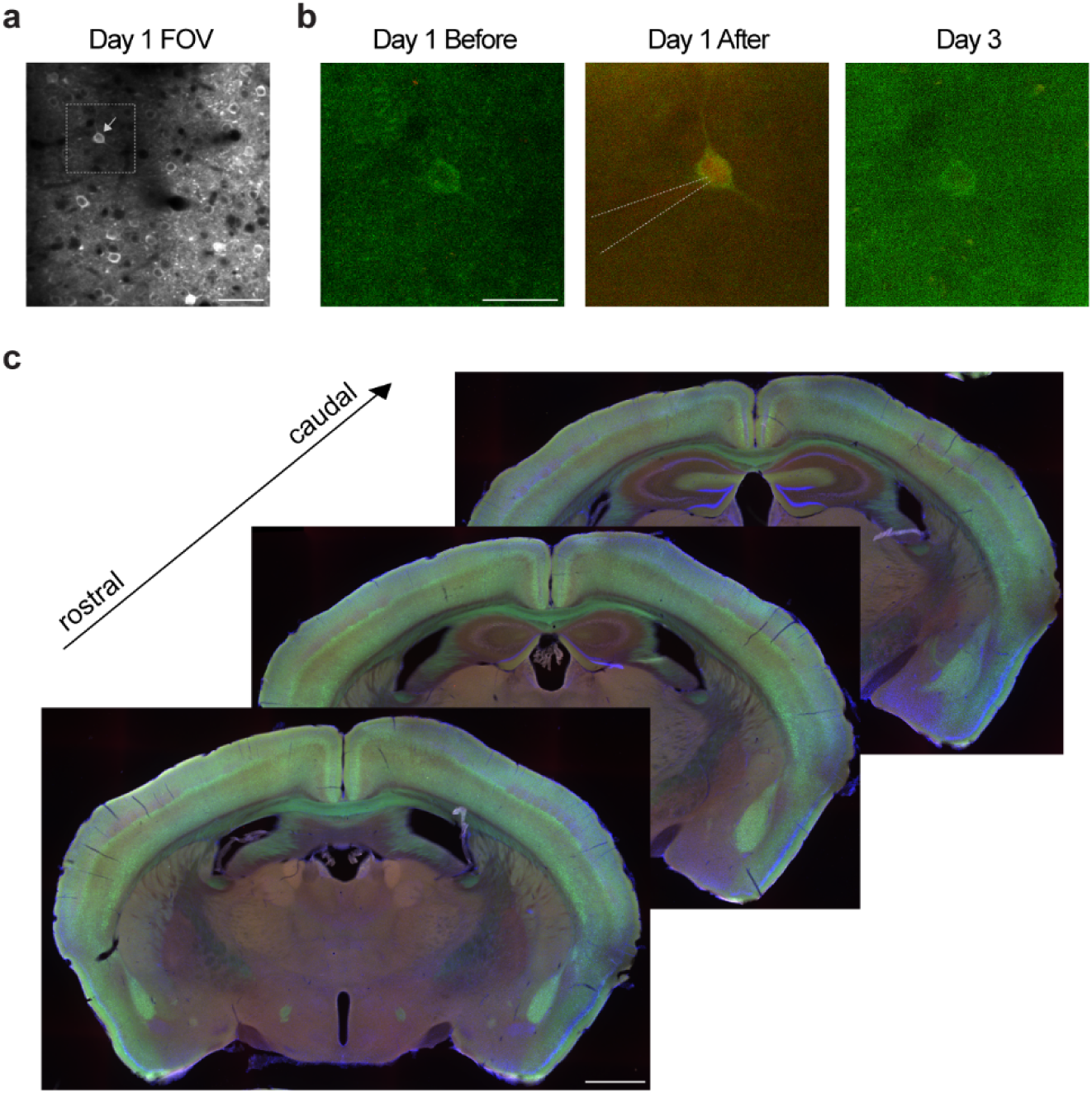
Absence of presynaptic labelling in sham single-cell based monosynaptic input tracing experiments. **a**, Imaging FOV for an example single-cell based monosynaptic retrograde tracing experiment. The arrow indicates a cell selected for electroporation based on its modulation during spontaneous movements. Scale bar, 50 µm. **b**, Day 1, target cell (as indicated in **a**) before and after 2-PT guided electroporation with Alexa594 and DNA (TVA, G, and mCherry). Note the entry of Alexa 594 into the target cell (red). Following electroporation, RV-RFP were injected close to the electroporated cell. Day 3, the target cell did not express the reporter protein mCherry, indicative of a failed transfection. Note that we used a single plasmid encoding for TVA, G, and mCherry. Scale bar, 25 µm. **c**, Histological analysis of the example experiment (**a**-**b**). Scale bar, 1 mm. We did not observe the emergence of a presynaptic cells (*n* = 4 brains). In a separate mouse, we implanted a custom cranial window, and we injected RV-RFP through the access port without electroporation to test off-target effect of RV (control injection). Also in this case, we did not observe the emergence of a presynaptic cells. Red, RFP. Green, GCaMP6s. Blue, NeuroTrace 435/455.

**Extended Data Fig. 9:**
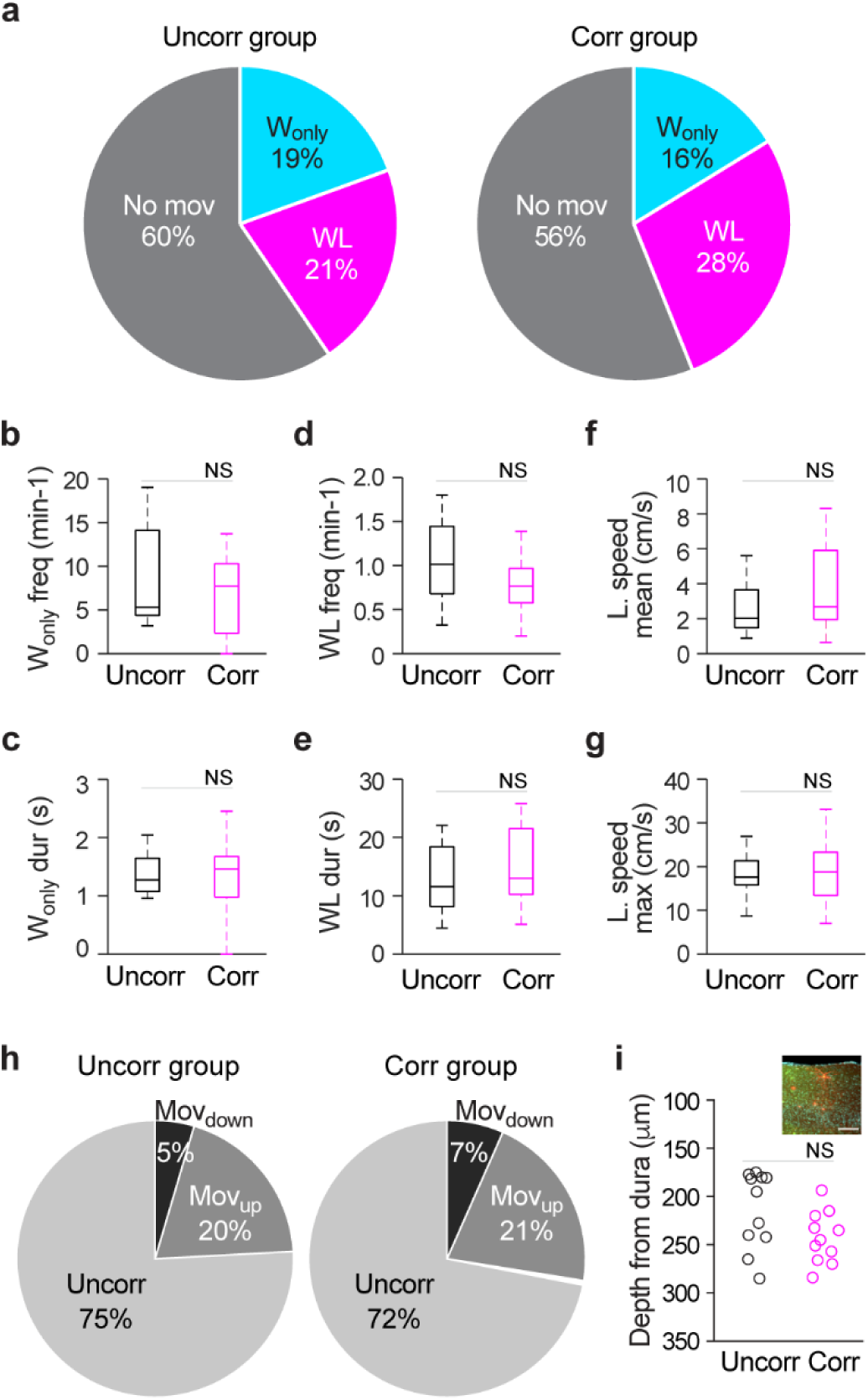
Spontaneous movements and neuronal properties of the movement-uncorrelated and movement-correlated groups for monosynaptic input tracing. **a**, Pie chart of time spent per spontaneous movement type (W_only_, WL) and not moving (*P* > 0.05 for movement-uncorrelated vs. movement-correlated, 2-way ANOVA, *n* = 11 per group). **b-g**, Spontaneous movement parameters, W_only_ frequency (**b**), W_only_ duration (**c**), WL frequency (**d**), WL duration (**e**), mean (**f**) and maximum (**g**) locomotion speed (*P* > 0.05 for all panels). **h**, Fraction of movement-uncorrelated, Mov_down_, and Mov_up_ neurons per group (*P* > 0.05 for uncorr. vs. corr., 2-way ANOVA). **i**, Cortical depth of each postsynaptic neuron. Depth was estimated starting from dura matter through in vivo 2-PT structural imaging (*P* > 0.05, Wilcoxon rank-sum test). Inset, epifluorescence image of a coronal brain slice encompassing wS1 and example postsynaptic neuron expressing both a red and green fluorescent protein (arrow). Red, RFP. Green, GCaMP6s/GFP. Cyan, DAPI. Scale bar, 100 µm.

**Extended Data Fig. 10:**
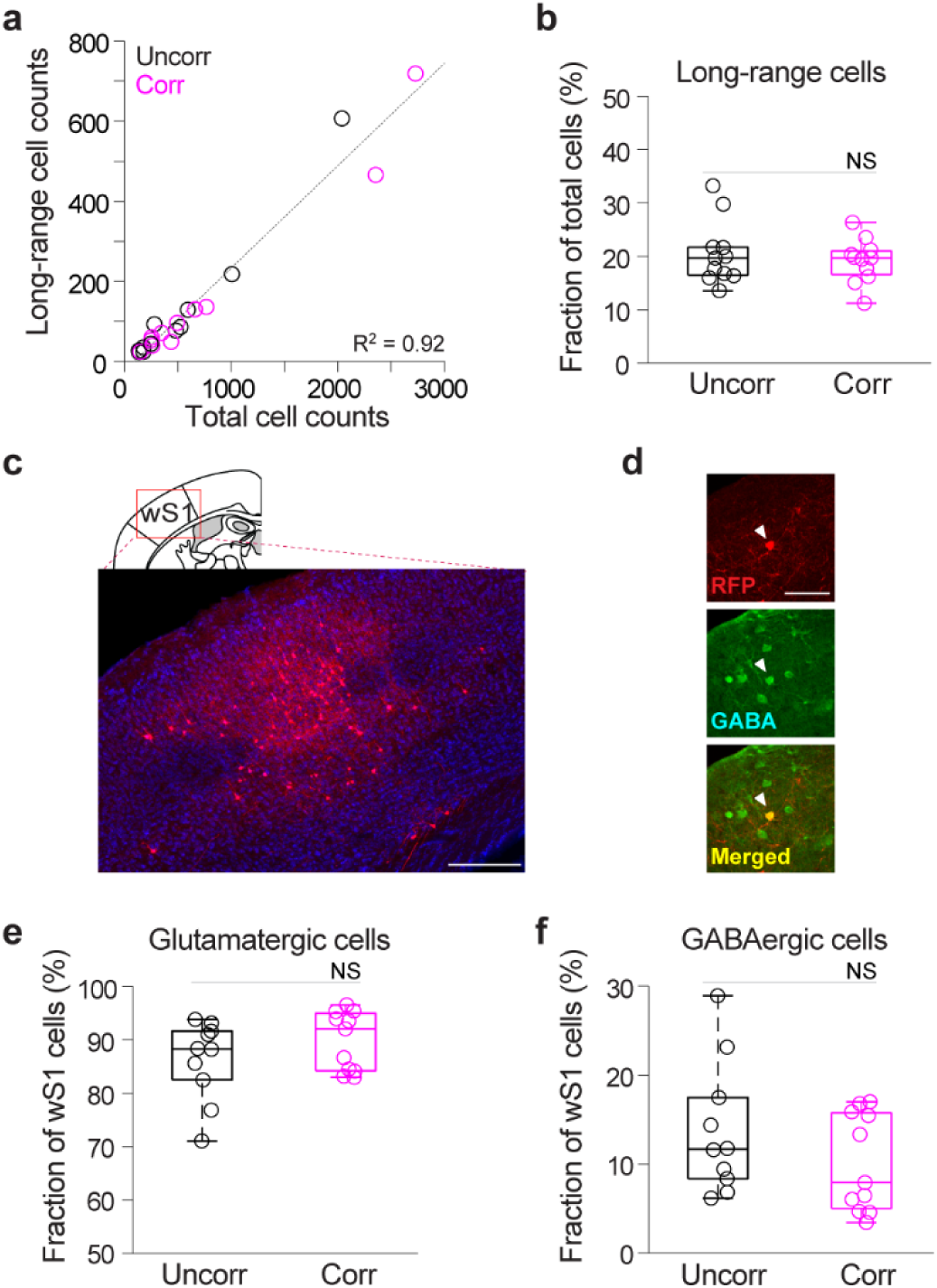
Long-range and local glutamatergic and GABAergic presynaptic neurons per brain. **a**, Number of long-range vs. total presynaptic neurons per brain (*P* < 0.001, regression). Black circles, movement-uncorrelated presynaptic neuron group (*n* = 11). Magenta circles, movement-correlated postsynaptic neuron group (*n* = 11). **b**, Long-range presynaptic neurons as fraction of total presynaptic neurons (*P* > 0.05, randomization test). **c**, Epifluorescence image of a coronal brain slice, wS1 (example presynaptic network included in Fig. 3a, d). Note the “hourglass” distribution of RFP^+^ presynaptic neurons across cortical layers 2/3, 4, and 5. Scale bar, 250 µm. **d**, Images denoting the co-localization of RFP (presynaptic neurons, Alexa 555, red) and GABA (GABAergic cells, Alexa 647, represented in green) in wS1. Scale bar, 50 µm. **e**-**f**, Glutamatergic (**e**) and GABAergic (**f**) presynaptic neurons as fraction of wS1 presynaptic neurons (*n* = 10 uncorr. and 11 corr., *P* > 0.05 for both panels, randomization tests).

**Extended Data Fig. 11:**
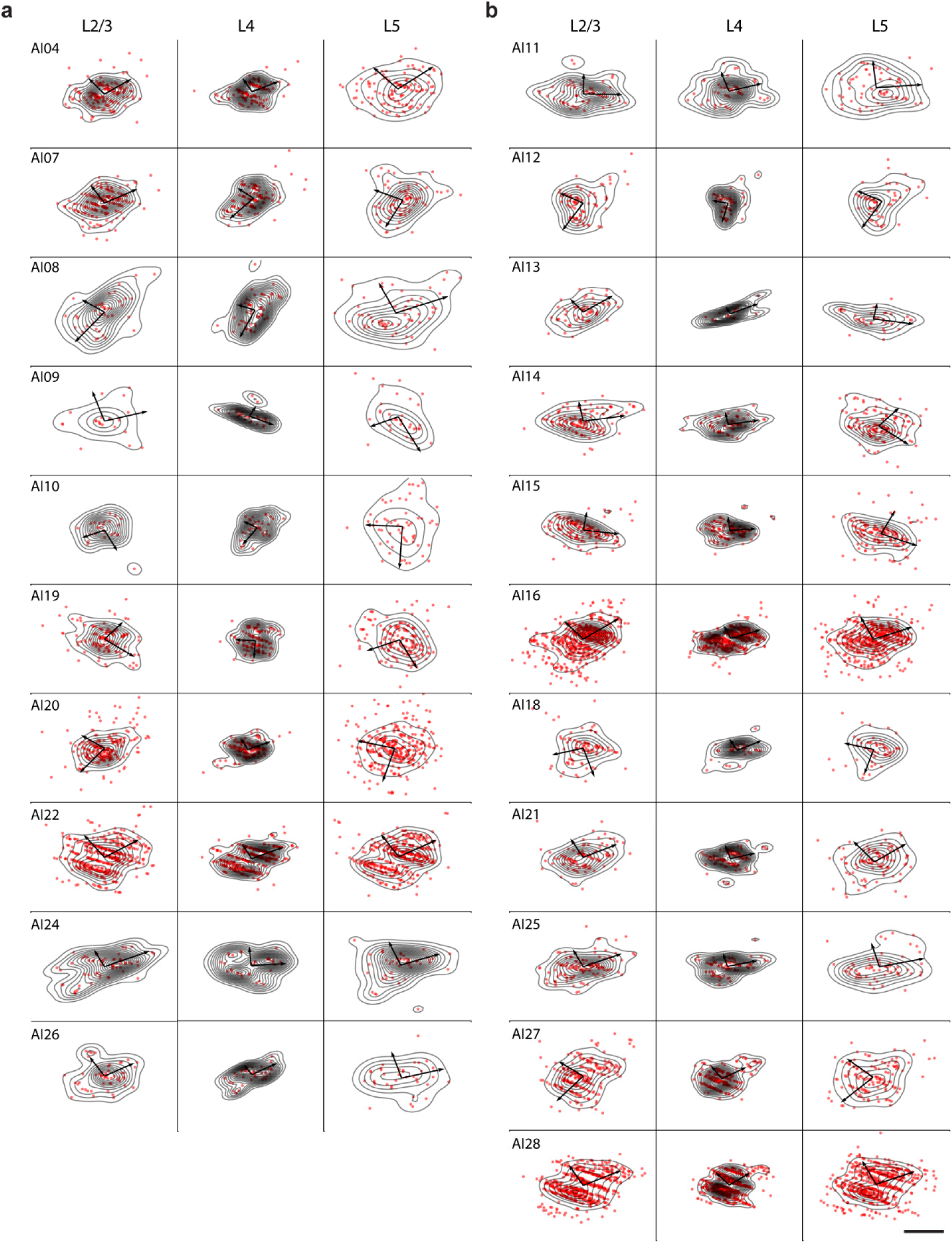
Spatial distribution of individual local glutamatergic presynaptic networks. Layer-by-layer horizontal flat projections of individual wS1 glutamatergic presynaptic networks with gaussian kernel Density estimation. We use the determinant of the estimated covariance as a measure of presynaptic networks dispersion. Smaller spatial dispersion of glutamatergic presynaptic cells in L4, than in L2/3 (*P* < 0.0001) and L5 (*P* < 0.0001) for all brains, independently of group (uncorr. and corr., one-sided *t*-test). **a**, Movement-uncorrelated postsynaptic group. **b**, Movement-correlated postsynaptic group. Scale bar, 500 µm.

**Extended Data Fig. 12:**
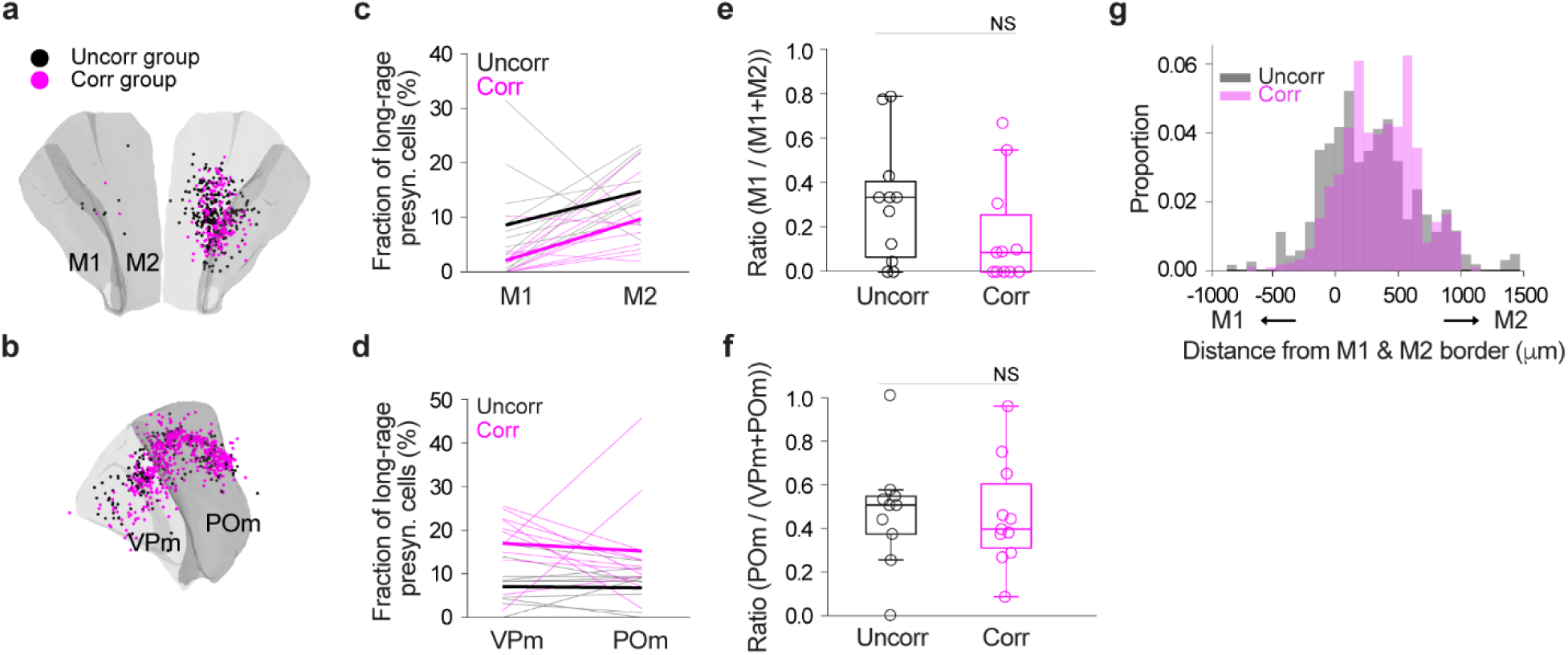
Motor cortical (M1 and M2) and thalamic (VPm and POm) presynaptic networks. **a**, Three-dimensional distribution of all presynaptic neurons within motor cortex (M1 and M2) for the movement-uncorrelated and movement-correlated postsynaptic neuron groups (data aligned to the Allen Mouse Common Coordinate Framework; uncorr. vs. corr.*, P* > 0.05, randomization test on 2-Wasserstein distance, *n* = 11 brains per group). **b,** As in **a**, but within sensory thalamic nuclei, VPm and POm. The spatial distribution of presynaptic neurons of across the sensory thalamus are similar between the two groups (*P* > 0.05, randomization test on 2-Wasserstein distance). **c,** M1 and M2 presynaptic neurons as fraction of long-range presynaptic neurons (uncorr. vs. corr.; M1, *P* < 0.05; M2, *P* = 0.057; Wilcoxon rank-sum test). Thin lines, individual data points. Thick lines, mean. **d,** VPm and POm presynaptic neurons as fraction of long-range presynaptic neurons (uncorr. vs. corr.; VPm, *P* < 0.01; POm, *P* < 0.05; Wilcoxon rank-sum test). **e,** Relative proportion of M1 vs. M2 neurons (*P* > 0.05, Wilcoxon rank-sum test). **f**, Relative proportion of POm vs. VPm neurons (*P* > 0.05, Wilcoxon rank-sum test). **g,** Weighted distribution of motor cortical presynaptic neurons as function of distance from the M1 and M2 border (uncorr. vs. corr., *P* > 0.05, randomization test).

**Extended Data Fig. 13:**
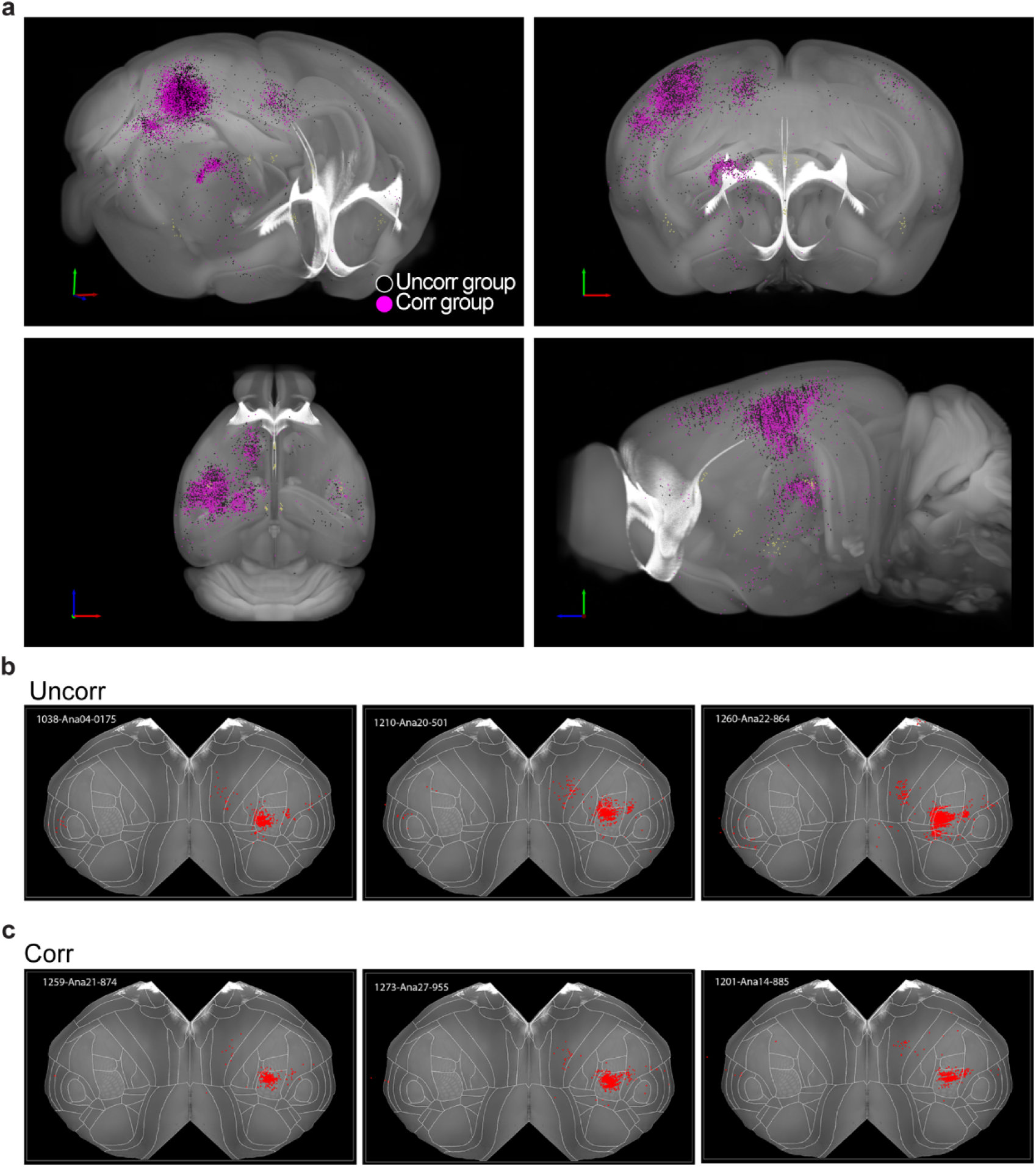
Brain-wide spatial distribution of presynaptic networks. **a**, Presynaptic networks of the movement-uncorrelated (*n* = 5801, 11 networks, black) and movement-correlated (*n* = 8699, 11 networks, magenta) groups superimposed on the Allen Mouse Common Coordinate Framework, denoting the high degree of spatial overlap of presynaptic neurons from both groups (*P* > 0.05 for each long-range area, randomization test on 2-Wasserstein distance). **b-c**, Cortical presynaptic networks from example movement-uncorrelated (*n* = 3, **b**) and movement-correlated (*n* = 3, **c**) brains (flat maps).

**Extended Data Fig. 14:**
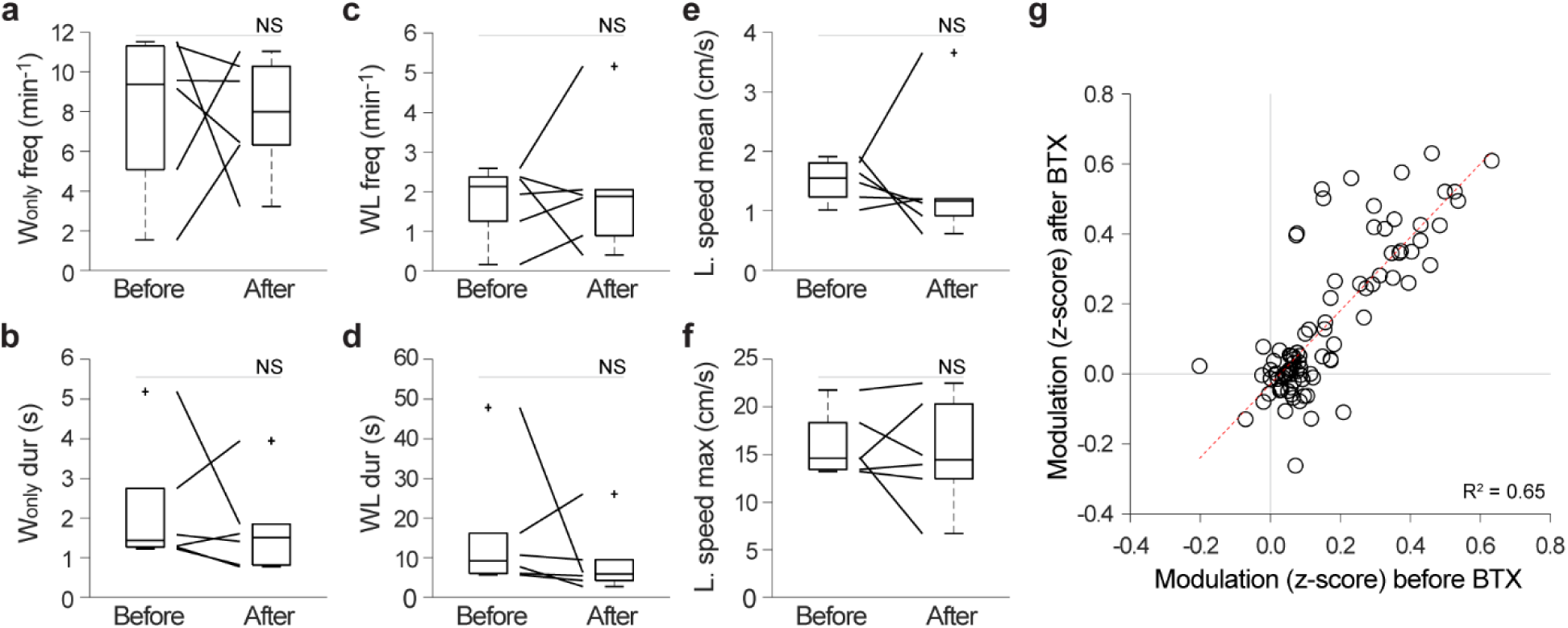
Neuronal activity and spontaneous movement before and after unilateral mystacial pad paralysis. **a-f**, Spontaneous movement parameters, W_only_ frequency (**a**), W_only_ duration (**b**), WL frequency (**c**), WL duration (**d**), mean (**e**) and maximum (**f**) locomotion speed before and after BTX injection in the mystacial pad (*P* > 0.05 for all panels, Wilcoxon signed-rank test, *n* = 6 FOVs, 2 sessions per FOV, 5 animals). **g**, Modulation of movement-correlated neurons during spontaneous movements (WL), before and after unilateral mystacial pad paralysis induced by BTX injection (*P* < 0.001, regression).

**Extended Data Fig. 15:**
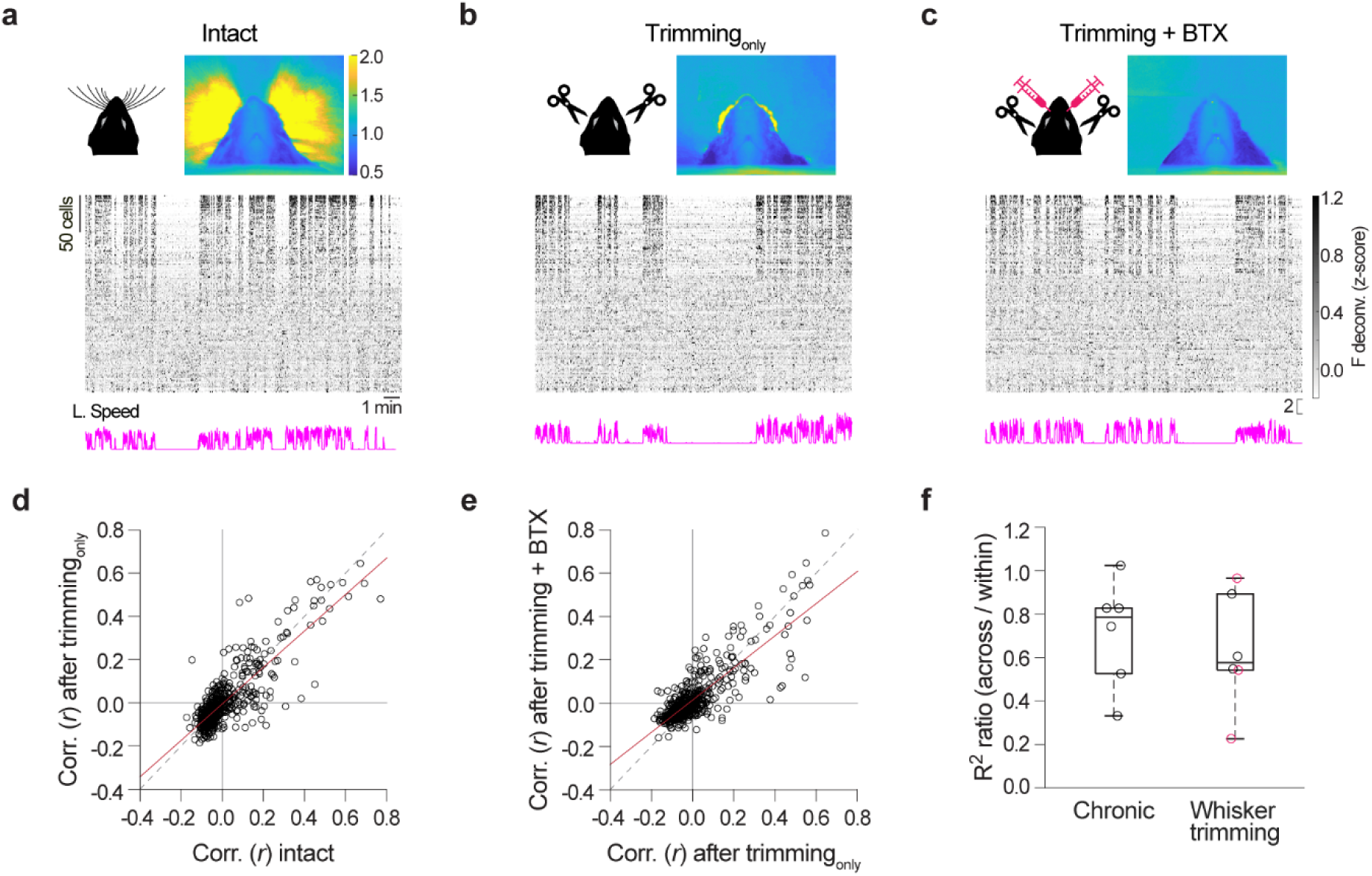
Effect of bilateral whisker trimming and whisker trimming combined with mystacial pad paralysis on wS1 L2/3 PNs activity in relation to spontaneous movements. **a-c**, Top, Example images of videorecorded whisker movements (mean of the absolute difference between consecutive frames over ∼14 s), before (intact, **a**) and after bilateral whisker trimming (Trimming_only_, **b**), as well as combined bilateral whisker trimming and bilateral BTX injections (Trimming + BTX, **c**) in the same mouse. Bottom, Raster plots of neuronal activity during spontaneous movements and corresponding locomotion speed traces; individual neurons (690) tracked across the three imaging sections (intact, Trimming_only_, and Trimming + BTX) are sorted from top to bottom by decreasing weight on the first principal component. **d**, Correlation (r) of the activity of individual neurons with locomotion speed before (intact) vs. after Trimming_only_ (R^2^ = 0.71, *P* < 0.0001, regression, *n* = 3 FOVs of 2 mice). **e**, Correlation (r) of the activity of individual neurons with locomotion speed Trimming_only_ vs. Trimming + BTX (R^2^ = 0.70, *P* < 0.0001, regression, *n* = 3 FOVs of 2 mice). **f**, Prediction of locomotion speed from population activity. Out-of-sample R^2^ ratio (R^2^_across_ / R^2^_within_) for chronic imaging (as in Fig. 1l), for intact vs. Trimming_only_ (black circles), and for Trimming_only_ vs. Trimming + BTX (magenta circles) (*P* > 0.05, Wilcoxon rank-sum test).

**Extended Data Fig. 16:**
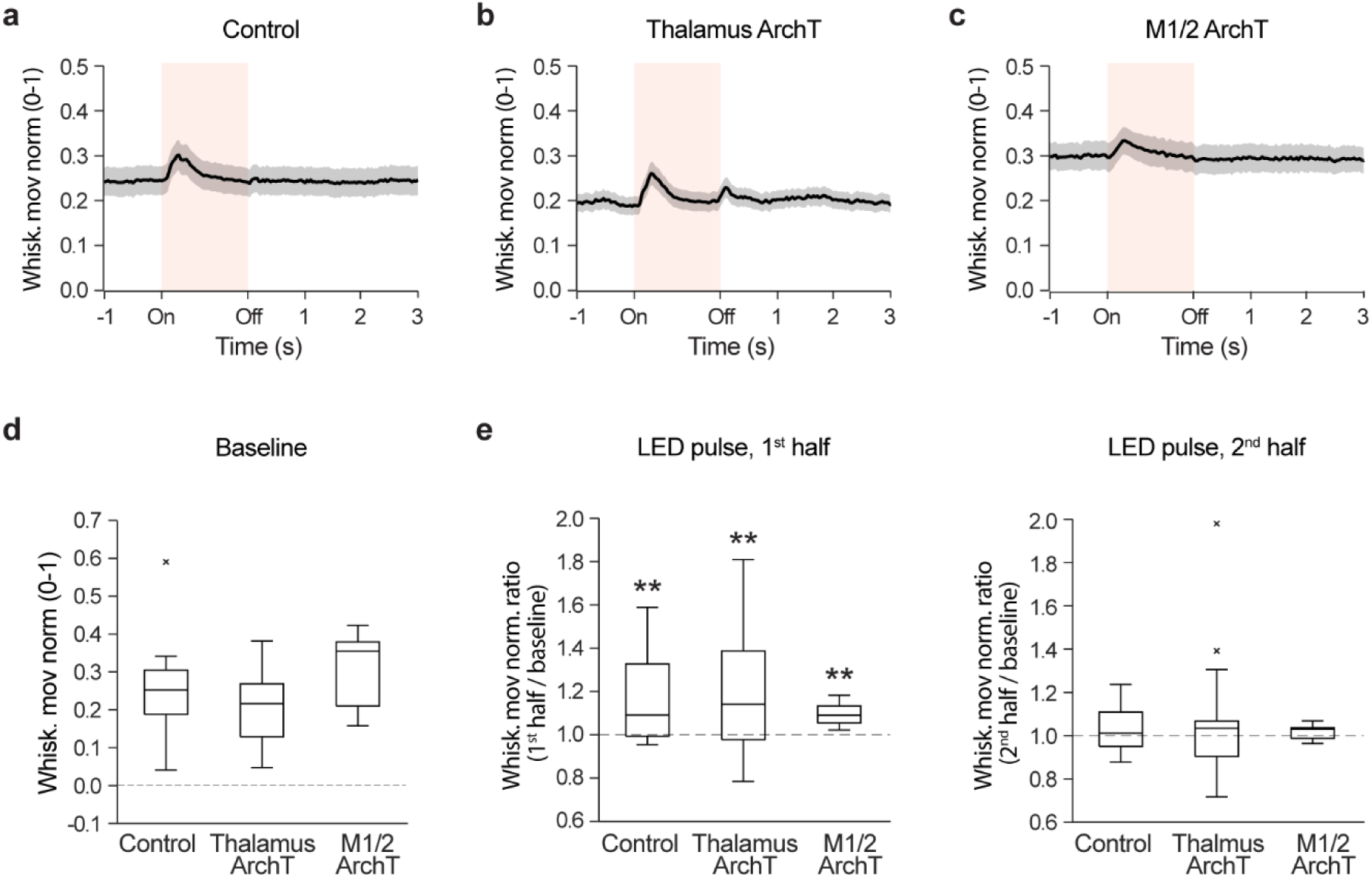
Behavioral effect of light presentation. **a-c**, Effect of light presentation on the spontaneous movements of control (*n* = 9, **a**), thalamus ArchT-expressing (*n* = 5, **b**), and M1/2 ArchT-expressing (*n* = 3, **c**) mice. Whisker movements aligned to the onset of light stimulation. **d**, Whisker movements during baseline (0.5 s window prior to light onset; *P* = 0.048, Kruskal-Wallis test across mouse groups). **e**, Quantification of behavioral changes over the first part (left, 0 - 0.5 s) and second part (right, 0.5 – 1/1.5 s) of the light pulse relative to baseline. All groups showed a brief (first half) increase in whisker movements locked to light presentation onset (*P* < 0.05, Wilcoxon signed-rank test whisker movements norm ≠ 1 vs. whisker movements norm = 1), and this increase was comparable across groups (*P* > 0.05, Kruskal-Wallis test). Whisker movements in the second half of the light pulse did not differ from that of baseline (*P* > 0.05, Wilcoxon signed-rank test whisker movements ratio ≠ 1 vs. whisker movements ratio = 1) and were similar across groups (*P* > 0.05, Kruskal-Wallis test).

**Extended Data Fig. 17:**
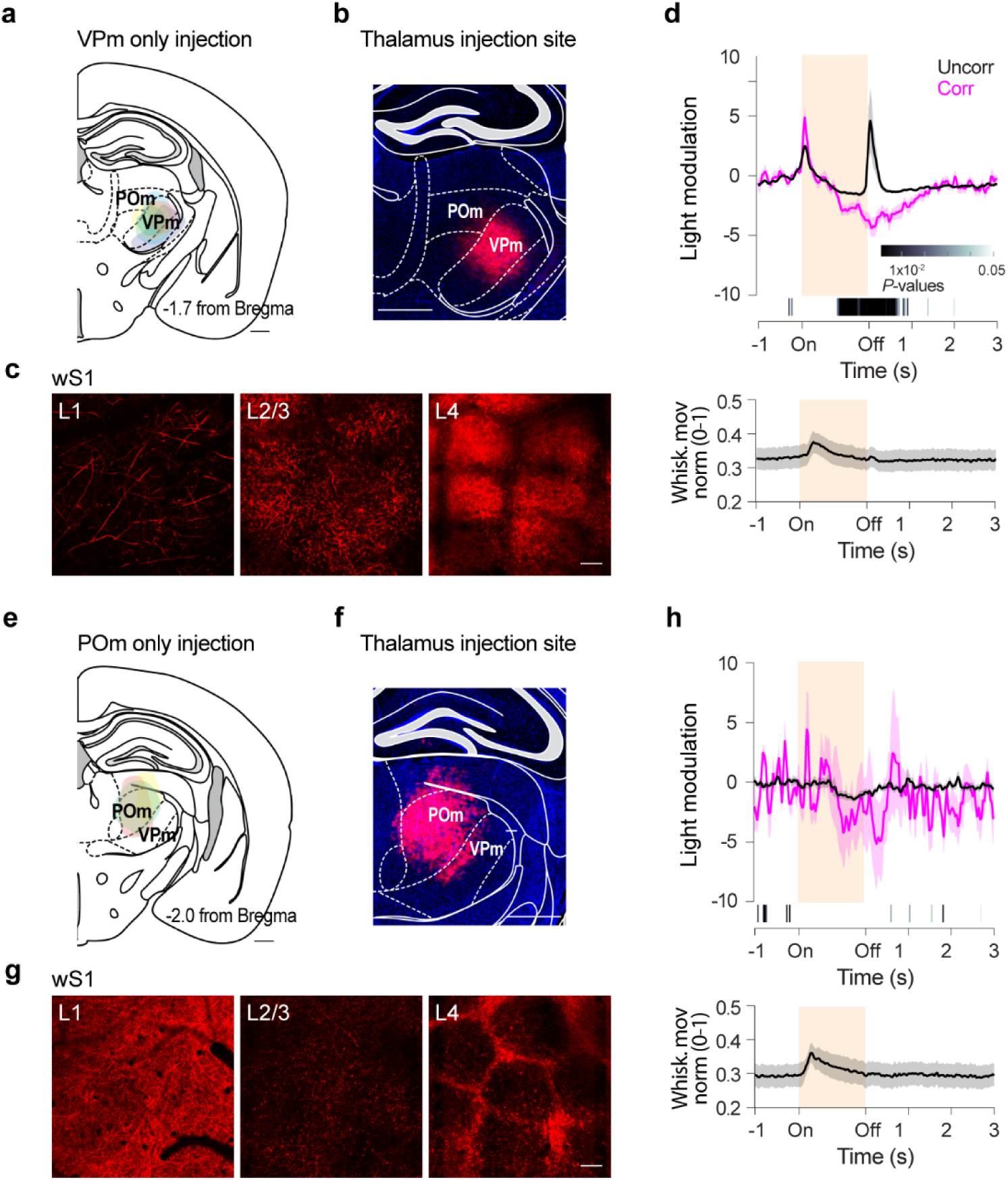
Optogenetic suppression of VPm and POm inputs in wS1 L2/3. **a**, Expression of the opsin ArchT in VPm. ArchT-tdTom^+^ areas are overlayed on the mouse atlas section corresponding to the center of the injection for VPm. Each shading color represents an individual mouse (*n* = 5). Scale bar, 0.5 mm. **b**, Example composite epifluorescence image of a coronal brain section showing ArchT-tdTom^+^ cell bodies in VPm. Scale bar, 0.5 mm. **c**, Laminar distribution of VPm ArchT-tdTom^+^ axons in wS1, as demonstrated by example 2-PT images. The characteristic patterns of innervation confirm the selectivity of injection during imaging, prior to histological analysis. Scale bar, 0.1 mm. **d**, Baseline-subtracted mean activity (light modulation) of movement-uncorrelated (black) or movement-correlated (magenta) neurons significantly affected by light pulses (*n* = 174 of 1108 uncorr., and *n* = 72 of 338 corr., 6 mice). *P* values, comparison of activity of movement-uncorrelated and movement-correlated neurons (Wilcoxon rank-sum test). Note that the movement-uncorrelated group includes 8 neurons (of 174) with a strong rebound activity; running analysis with and without these neurons did not affect the results. **e-h**, As in a-d, but for the expression of the opsin ArchT in POm (neurons significantly changed by light stimulation, *n* = 32 of 585 uncorr., and *n* = 17 of 267 corr., 3 mice).

**Extended Data Fig. 18:**
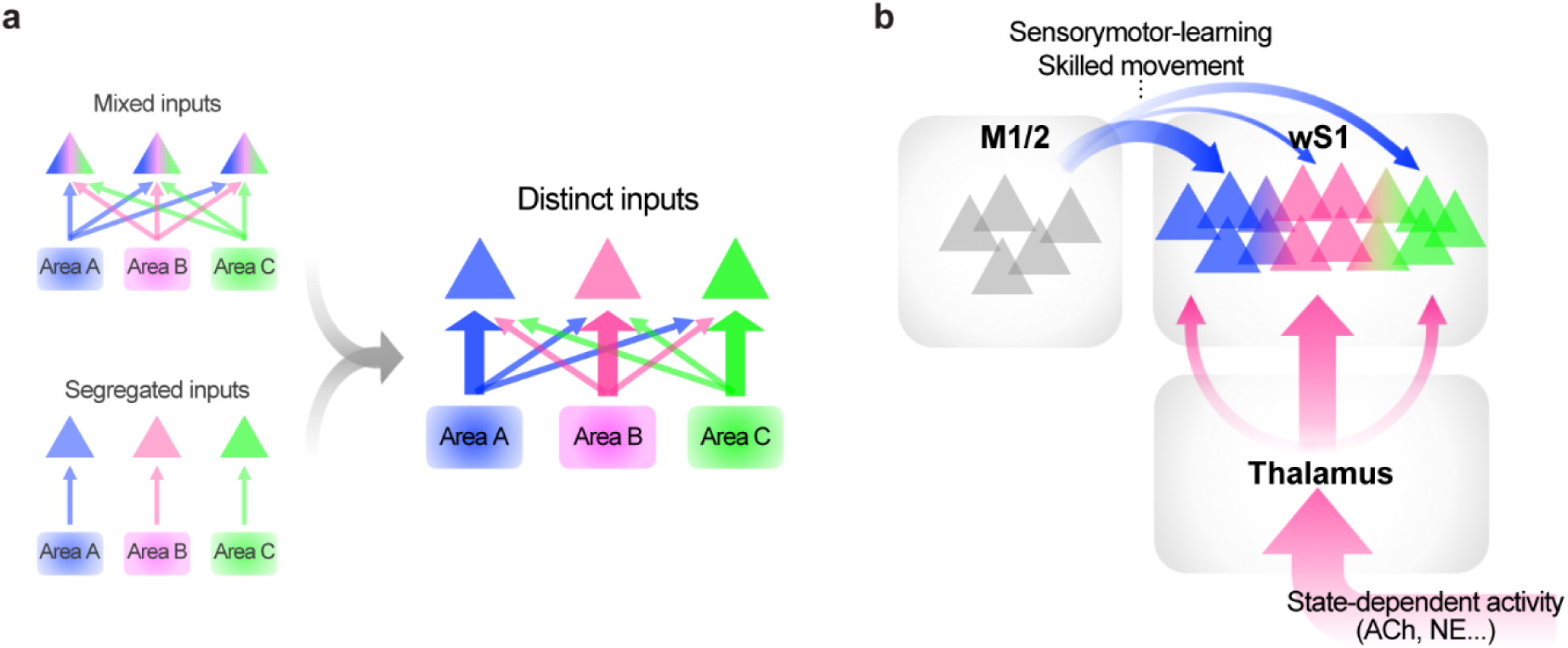
Summary of the anatomical presynaptic connectivity rule and its proposed functional implications. **a**, Presynaptic connectivity rule underlying the functional heterogeneity of cortical PNs. Movement-uncorrelated and movement-correlated neurons did not receive separate sets of inputs nor completely random or mixed inputs from the distinct wS1-projecting areas. Instead, movement-uncorrelated and movement-correlated neurons had distinct presynaptic networks characterized by a selective fractional decrease in M1/2 inputs and increase in thalamic inputs. **b**, Hypothetical functional significance of distinct long-range inputs to wS1. While thalamic inputs may convey behavioral state-dependent activity, the M1/2 projection may instead be required for more complex changes in wS1 activity structure, such as establishment of new sensorimotor associations or acquisition of skilled movements.

## Notes

### Competing Interest Statement

The authors have declared no competing interest.

### Summary of Updates

Updated text and figures, and new results in: Fig. 2 (updated); Fig. 5 (updated); Extended Data Fig. 4 (new); Extended Data Fig. 12 (updated; previously Extended Data Fig. 11); Extended Data Fig. 13 (updated; previously Extended Data Fig. 12); Extended Data Fig. 15 (new); and Extended Data Fig. 17 (new).

